# Open Educational Resources for distributed hands-on teaching in molecular biology

**DOI:** 10.1101/2024.03.28.587173

**Authors:** Ariel Cerda, Alejandro Aravena, Valentina Zapata, Anibal Arce, Wladimir Araya, Domingo Gallardo, Javiera Aviles, Francisco Quero, Isaac Nuñez, Tamara Matute, Felipe Navarro, Valentina Ferrando, Marta Blanco, Sebastian Velozo, Sebastian Rodriguez, Sebastian Aguilera, Francisco Chateau, Ruby Olivares-Donoso, Jennifer C Molloy, Guy Aidelberg, Ariel B. Lindner, Fernando Castro, Pablo Cremades, Cesar Ramirez-Sarmiento, Fernan Federici

## Abstract

The urgent need to develop a more equitable bioeconomy has positioned biotechnology capacity building at the forefront of international priorities. However, in many educational institutions, particularly in low-and middle-income countries, this remains a major challenge due to limited access to reagents, equipment, and technical documentation. In this work, we describe Open Educational Resources (OER) composed of locally produced biological reagents, open source hardware and free software to teach fundamental techniques in biotechnology such as LAMP DNA amplification, RT-PCR RNA detection, enzyme kinetics and fluorescence imaging. The use of locally produced reagents and devices reduces costs by up to one order of magnitude. During the pandemic lockdowns, these tools were distributed nationwide to students’ homes as a lab-in-a-box for remote teaching of molecular biology. To test their performance, a total of 93 undergraduate students tested these resources during a biochemistry practical course. 27 out of 31 groups (∼87%) successfully achieved the objectives of the PCR activity, while 28 out of 31 groups (∼90%) correctly identified the target using LAMP reactions. To assess the potential application in secondary school, we organized three workshops for high school teachers from different institutions across Chile and performed an anonymous questionnaire, obtaining a strong agreement on how these OER broaden teachers’ perspectives on the techniques and facilitate the teaching of molecular biology topics. This effort was made possible through a close collaboration with open source technology advocates and members of DIYbio communities, whose work is paving the way for low-cost training in biology. All the protocols and design files are available under open source licenses.

## Introduction

Biology has been declared as a discipline of central importance for the 21st century. The pressing need to develop more sustainable materials, healthier food and cleaner energy has positioned biotechnology at the forefront of international priorities. The COVID pandemic has also evidenced the urgent need for more evenly distributed capacities in biotechnology to ensure timely responses to public health crises. However, building capacity in biotechnology remains a major challenge due to limited access to reagents, instruments, and technical documentation in many educational institutions around the world. These barriers hinder training the next generation of scientists and limit opportunities to spark the student’s interest in biological sciences and biotechnologies. Developing Open Educational Resources (OER) tailored to address these challenges is therefore essential to support equitable access to biotechnology education. While recent developments in synthetic biology are expanding the possibilities for hands-on training in life sciences [1–3], capacity building in biotechnology remains particularly difficult in low-and middle-income countries (LMICs), where access to essential instruments and reagents is often limited. Reagents are imported at high costs and scientific equipment, such as fluorescence microscopes and thermocyclers, is either unavailable or inaccessible because of the high demand for research. The establishment of scientific infrastructure solely dedicated to teaching remains a scarce luxury for most Latin American universities—and very uncommon in high schools.

In response, several researchers are turning to open source hardware as a solution for this challenge [4,5]. The progress on frugal biotechnology [6,7], DIYbio movements [8–10], standardized genetic resources [11], initiatives on open scientific hardware [12], and ip-free resources for the local production of reagents [13–15] have collectively paved the way for a more equitable access to hands-on training in molecular biology. These resources can enable personalized environments to learn instrument operation, hands-on laboratory practices and experimental design, which are a vital part of life science education for both theory reinforcement and practical skills development.

In this work, we describe the development of Open Educational Resources for low-cost, hands-on teaching in molecular biology that integrate advances in open source technology. We implemented a lab-in-a-box that included all the necessary reagents and materials for students to gain training in molecular biology techniques at home such as RT-PCR, DNA amplification, fluorescence and enzyme kinetics. These resources consist of locally produced reagents, open source hardware as well as free software and collaborative notebooks. All the protocols and design files are available under open source licenses that explicitly grant permission for study, replication and commercialization of the resources. We also describe how these resources were used for remote teaching during the COVID-19 lockdowns, including their implementation in molecular biology practicals performed by 93 third-year biochemistry students during 2020 and 2021, as well as capacity training of 37 high school teachers in Chile. To evaluate their educational potential and limitations, we conducted an online questionnaire to high school teachers trained in the use of these OERs. This study aims to assess the feasibility, accessibility, and pedagogical effectiveness of these OER-based molecular biology kits in education.

## Results

### RT-PCR for the detection of SARS-CoV-2

The first practical consisted of the reverse transcription polymerase chain reaction (RT-PCR), a technique widely used for converting RNA into DNA and amplifying specific sequences, making it the gold standard for detecting RNA viruses in molecular diagnostics [16–18]. We focused on detecting the Severe Acute Respiratory Syndrome Coronavirus 2 (SARS-CoV-2) for two main reasons: first, the students’ motivation for learning about the virus causing the COVID19 pandemic; and second, the availability of low-cost and open-access RT-PCR reagents for SARS-CoV-2 detection, a result of ongoing local initiatives [19–22]. Despite the relevance of this technique on molecular diagnostics, its broader implementation in teaching labs, especially in developing countries, has been limited by the high costs and scarce availability of necessary reagents and equipment. By the end of this practical, students were expected to perform RT-PCR to detect synthetic SARS-CoV-2 RNA, understand the principles of RNA handling and amplification, and interpret gel electrophoresis results.

We integrated two open-source projects for a cost-effective solution: the PocketPCR and off-patent enzymes. The PocketPCR is a compact, open-source USB-powered thermocycler designed to make PCR technology more affordable and portable [23](**Fig 1 and S1 Fig**). This open hardware device facilitates PCR reactions through precise, easily adjustable temperature cycles, and can be powered by a standard USB-C charger or computer, making it ideal for use outside traditional laboratory settings. The pocketPCR can be fabricated from online blueprints or acquired commercially for 100 € from GaudiLabs. The DNA coding for the open enzymes, on the other hand, were freely sourced from the Reclone network and FreeGenes project [13,24], which facilitates open access to DNA plasmids encoding for public domain diagnostic enzymes [19–22,25]. For instance, the FreeGenes project and ReClone network have engaged researchers around the globe who share DNA plasmids encoding these off-patent enzymes and protocols for their local production. These collaborative efforts have enabled many laboratories to access molecular diagnostics at a reduced cost.

**Figure 1:**
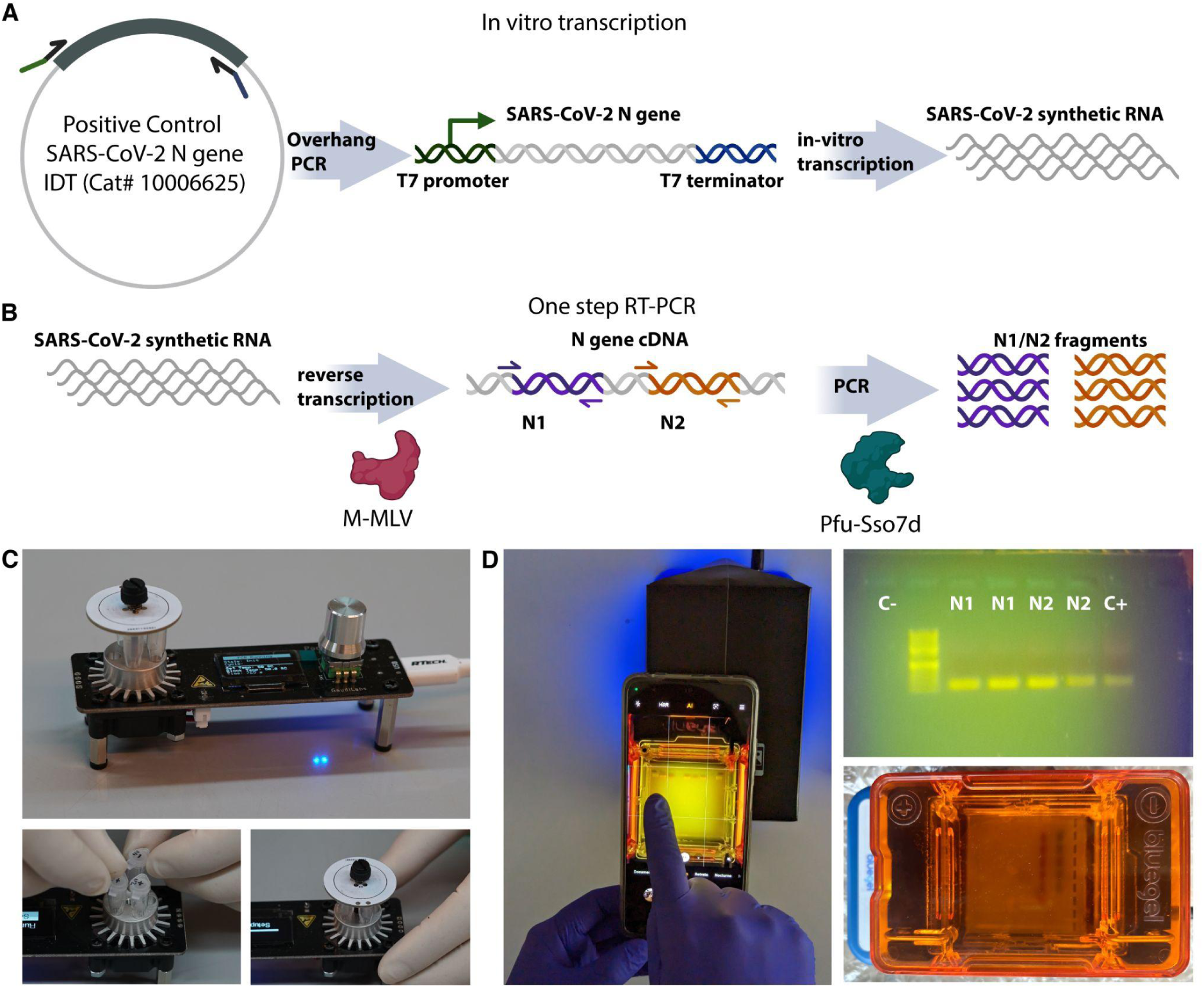
Schematic representation of RT-PCR practical. a) Positive control RNA produced by *in vitro* T7 RNAP transcription from a linear PCR product containing a T7 promoter and terminator. The PCR-amplified linear template was obtained from a plasmid harboring the “Positive Control SARS-CoV-2 N gene” from IDT (Cat# 10006625). b) Schematic representation of RT-PCR reaction performed during the practical. c) PocketPCR device holding up to five tubes. d) BlueGel™ electrophoresis system showing RT-PCR practical results (top right).

To design an experiment that emulates a diagnostic procedure for SARS-CoV-2, we used synthetic RNA that corresponds to the full-length Nucleocapsid (N) gene of approximately 1260 bp of SARS-CoV-2 (**Fig 1A**). The synthetic RNA was generated by *in vitro* transcription from a PCR-amplified linear DNA template obtained with custom-designed primers containing a T7 promoter and a terminator [19,20](**Fig 1A**). Additionally, we selected primers used in clinical diagnostics that target different regions of the SARS-CoV-2 N gene (N1 and N2, **Fig 1B**). The RT-PCR reaction was performed in a PocketPCR device (**Fig 1C**) using a master mix containing recombinant M-MLV reverse transcriptase and Pfu-Sso7d polymerase, produced locally [20]. The cost of the homemade reaction is approximately one order of magnitude lower than that of the commercial version (approximately $0.80 per reaction compared to $8.00 for the commercial kit). The protocols used for the local production of these enzymes are openly accessible at protocols.io [26,27]. In this practical, each student was provided with purified enzymes (M-MLV-RT and Pfu-Sso7d), reaction buffer, primers N1 and N2, a cDNA sample control of SARS-CoV-2 N gene, and two RNA samples (“positive” and “negative” for the presence of the synthetic viral RNA fragment), transported and stored at room temperature. To avoid the use of personal refrigerators, the RNA samples, buffers and enzymes were kept at room temperature for the ten days of transit among the three students in each group. Given the lack of real-time reading capabilities, we avoided the quantitative aspects of the diagnostic process and focused on a qualitative assay, aiming solely for the detection of the presence or absence of synthetic viral RNA verified by the presence of an amplification band on a gel. Although open source and DIY electrophoresis chambers have been developed, we opted for the BlueGel® device that has been CE certified (Amplyus, MA, USA) (**Fig 1D**). This device has an integrated blue LED array for visualization and it has been used for SARS-Cov-2 detection [28].

As explained above, the experiments conducted by students were carried out over two consecutive years. During 2020, 11 out of 15 groups (∼73%) achieved the objectives of the activity. A group was considered successful if at least one of the three members was able to distinguish the positive sample from the negative one using the provided enzymes and reagents. The primary challenges encountered included partial evaporation of reagents during transportation, student inexperience in loading agarose gels that often led to wells being punctured, and potential degradation of the synthetic RNA. Most of these issues were resolved in the following year, when 100% of all 16 groups successfully met the activity objectives (a representative figure of this group’s results is shown in **Fig 1D**). While this experimental setup does not exactly replicate the conditions encountered in clinical diagnostics, it offers a valuable approximation on the fundamental principles of RNA handling, amplification and analysis. Students were also trained in primer design and applied this knowledge to understand results of this practical.

### LAMP reactions for DNA detection

The second practical focused on DNA amplification via loop-mediated isothermal amplification (LAMP). Although RT-PCR remains the gold standard in molecular diagnostics, alternative nucleic acid amplification methods have proliferated in the past two decades to provide faster, simpler, and more portable reactions [29]. Among these methods, LAMP (loop-mediated isothermal amplification) has gained popularity due to the use of a polymerase with strand-displacement activity that enables amplification at a constant temperature without the need for thermal cycling [30]. The isothermal nature of this reaction, along with a higher tolerance to reaction inhibitors, makes LAMP reactions attractive for point-of-care applications in resource-limited settings lacking specialized instruments and unprocessed samples. These features have put LAMP at the forefront of molecular developments for nucleic acid amplification in the context of point-of-care diagnostics. Through this practical, students were expected to apply the LAMP technique to detect transgenic DNA, prepare and analyze food samples, and use custom-built devices for result visualization. Additionally, they were introduced to programming and data analysis using Google Colaboratory notebooks, enabling them to interpret their findings while discussing the broader implications of GMOs.

Due to the robustness of LAMP, it is feasible to train students on applying this technique to unprocessed samples available at home [31]. We applied the *GMO Detective* project, an open citizen-science and educational endeavor towards an informed perception of GMOs [32], to detect transgenic DNA from vegetables and processed foods (e.g. veggie burger). *GMO Detective* consists of LAMP reactions that detect the presence of the CaMV 35S promoter, a commonly used sequence in the generation of transgenic plants. This kit applies the Quenching of Unincorporated Amplification Signal Reporters (QUASR) approach for end point determination [33], consisting of a fluorescently-labeled primer and a quencher-bearing probe. Positive LAMP QUASR reactions exhibit a fluorescent signal when the primer is incorporated into the amplified product and no signal when the probe is quenched (no amplification or no specific amplification). Sample preparation was simple, consisting of grinding and heating a small vegetable or processed food piece (**Fig 2A**).

**Figure 2:**
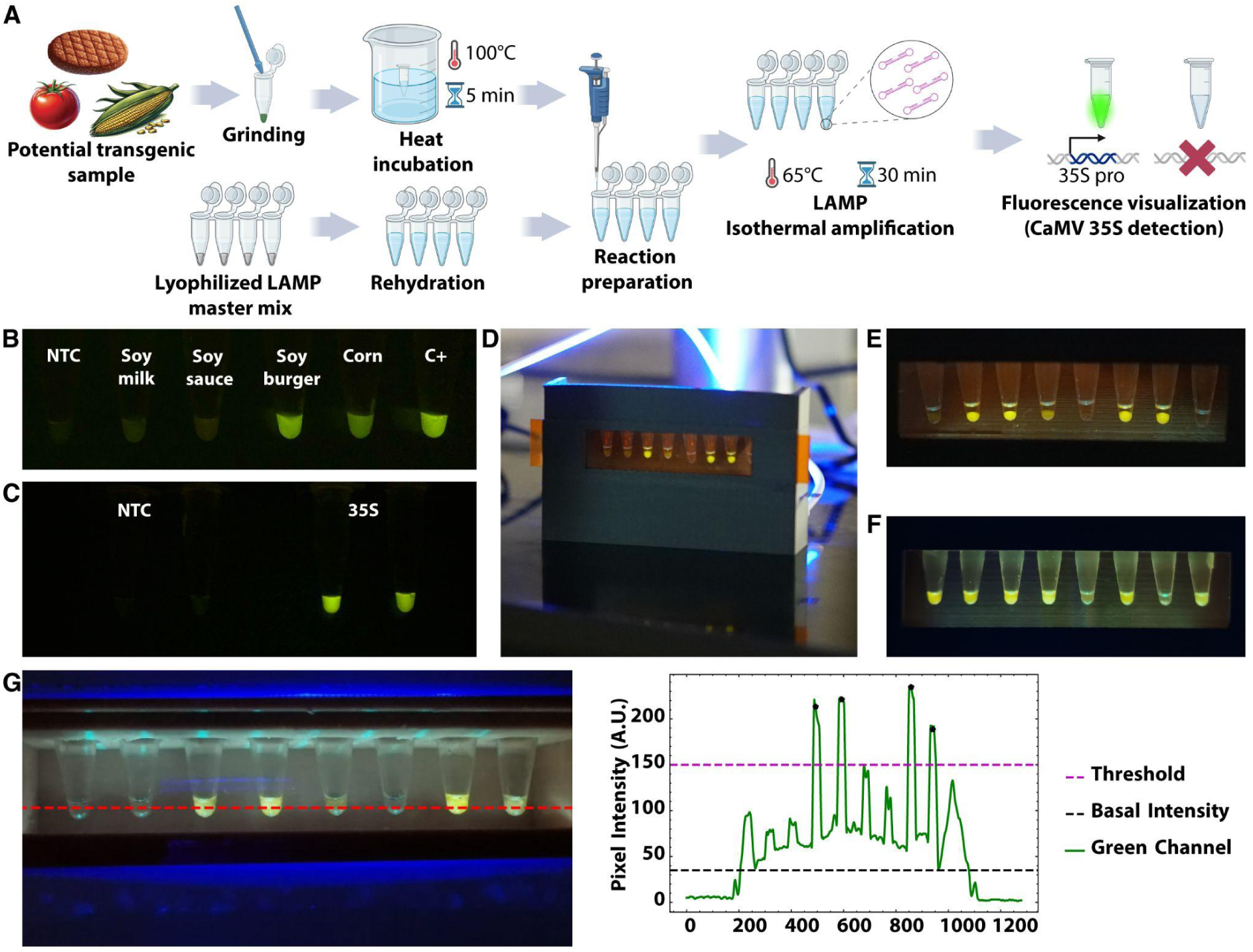
GMO Detective practical. a) Schematic representation of GMO Detective practical. b) Detection of 35S promoter in different soy-based foods. c) Reaction performance of freeze-dried reactions after two months storage at room temperature (two replicates are shown). d) Image of the 3D printed device being used by students. e) and f) Comparison of reaction with (e) and without (f) mineral oil. Mineral oil was added to avoid reaction evaporation and subsequent condensation on the tube lids. g) Image processing and analysis performed using Jupyter Notebooks designed for this course (Image processing and analysis code available from [34]) and run on Google Colaboratory. The red line indicates the position at which intensities were quantified. All images are raw images without editing, except for panel b and c, for which linear exposure was enhanced across the whole image for better dynamic range.

The first *GMO Detective* reactions were prepared in France using commercial reagents, lyophilized, and then shipped to Chile. However, this is unsustainable due to the high costs of shipping and delays in importation. Consequently, local production of *GMO Detective* reactions appears as a more suitable solution. Yet, this approach is limited by the high cost of commercial *Bst* polymerase, which must be imported under cold-chain conditions from the northern hemisphere. Additionally, the presence of glycerol in commercial enzyme buffers hinders lyophilization [35]. To overcome this barrier, we developed a local version of *GMO Detective* using homemade reagents and an off-patent *BstLF* polymerase from the openly distributed FreeGenes collection [13,36,37] (**S1 Appendix**). The costs of the homemade reactions is USD 0.36 per reaction, one order of magnitude cheaper than the costs of reactions implemented with commercial enzymes (In Chile, the NEB WarmStart kit costs USD 3.86 per reaction while NEB RTx and Bst2.0 WarmStart enzymes bought separately cost USD 3.38 per reaction). Before integrating them into the course, we validated these locally produced reactions using a range of soy-based food products likely to contain GMO soy, along with synthetic positive controls (**Fig 2B**). These locally produced LAMP reactions can be prepared with glycerol-free enzymes, lyophilized and stored at room temperature without any apparent loss in activity [36,38](**Fig 2C**). Freeze-drying the reactions without magnesium or reaction buffer—keeping them in liquid format until rehydration—allows for at least two months of storage at room temperature or 4°C (**S2 Fig**). Conversely, including them in the lyophilization step led to signal recovery after two months only if the lyophilized tubes are stored at 4 °C (**S2 Fig**). We also produced 35S and COX1 positive control vectors and tested an alternative quencher that is more affordable in Chile (**S1 Appendix**). Additionally, we tested a Texas Red fluorophore (red fluorescence) for the COX1 primer set, enabling dual color QUASR LAMP when used with FAM-labeled 35S primer set (**S1 Appendix**). The end-point results of these LAMP reactions can be monitored with a low-cost device composed of laser-cut acrylic pieces, a custom-made PCB, low-cost lee filters and 5 mm blue LEDs (emitting at 470 nm) [39]. To scale up the number of devices used in this practical, we created a 3D-printed version of the visualization device (**Fig 2D and S3 Fig**), further optimized by other groups (e.g. UROS project at Hackteria). Details and OpenSCAD editable design files are provided in [39]. We also replaced the original *GMO Detective* blue LEDs for 5 mm blue LEDs from SuperBright LEDs (Forward current: 30mA @ 3.3V, Peak forward current: 100mA, Max voltage forward: 3.8V), already used in the IORodeo blue transilluminator [40] and FluoPi fluorescence imaging device [41] for green fluorescence.

The practical began with the preparation of test samples (**Fig 2A**). For this, food samples, such as soy burgers, were mixed with nuclease-free water and ground in a 1.5 ml tube. These samples were incubated for 5 minutes in water heated to 95-100°C (**S4 Fig**). *GMO Detective* LAMP reactions were performed in standard PCR tubes incubated at 63°C for 35 min, followed by a 15-minute cooling period at room temperature before visualizing the results using a 3D-printed device (**Fig 2D-F**). To prevent evaporation and condensation, mineral oil was added to the reaction tubes (**Fig 2E and F**). Reactions were carried out with two technical replicates and repeated in triplicate on different days and at different students’ homes. For the 63-65°C incubation, students used commercially available 12V electronic coffee mugs [41,42] or sous-vide heaters [43] to create a temperature-controlled water bath. In some cases, students mixed boiling and cold water to achieve the desired temperature, monitored with a digital thermometer (**S4 Fig**). This water bath was prepared within the small plastic box provided in the lab-in-a-box (**S4 Fig**). Care was taken to properly seal the tubes to prevent water infiltration. Notably, the results were consistent across different heating methods, including the PocketPCR. Each student was guided to assemble an LED visualization device, prepare the food sample, conduct the reactions using different water-bath approaches, and report their findings. In both 2020 and 2021, students were provided with food samples, one transgenic and the other non-transgenic, with the objective of identifying each using the supplied reagents. The activity was successful in both years, with 14 of 15 groups in 2020 (93%) and 14 of 16 groups in 2021 (88%) correctly identifying the samples. The most common issues encountered were related to DNA extraction failures from the tissue samples (evidenced by the lack of COX control amplification) or contamination of the negative controls, leading to inconclusive results regarding sample identity. Representative results of their experiments can be found in **Figure 2E-F**. To perform quantitative analysis and discrimination of positive from negative reactions across the three replicates, students were provided with Google Colaboratory Jupyter notebooks designed to introduce students to basic coding, data analysis, and result visualization through shareable, cloud-based documents. The provided notebooks (Image processing and analysis; Fluorescence: Image processing and analysis from [34]) can be executed directly on Google Colaboratory and can be edited upon need, allowing students to analyze images taken with their phones and prepare plots for presentations and reports (**Fig 2G**).

Detecting GMOs in food samples offers a safer alternative to pathogen-focused diagnostics while actively engaging students in applying molecular biology concepts to investigate food composition. This approach serves as a springboard for learning about and debating the implications of GMOs in their daily lives.

### Basics of fluorescence readings

The goal of this section was to teach fluorescence measurement from biological samples. Particularly, how to appropriately select the best filter combination and illumination wavelengths to obtain high signal-to-background images of different fluorophores. This is a critical task encountered not only in microscopy but also in various other techniques (e.g. plate fluorometry, cell cytometry), hence, the use of fluorescence microscopes is not strictly necessary. Therefore, we designed a simple open hardware device called RGBfluor that captures the essence of fluorescence measurement without the complications of using an expensive and highly demanded fluorescence microscope. RGBfluor is a 3D printed device that consists of an illumination module controlled by a smartphone through Wi-Fi, a sample holder, and a multiple filter slider (**Fig 3A**). The illumination module consists of a custom-made printed circuit board (PCB) featuring ws2812 RGB LEDs and designed as a shield for the Lolin commercial board based on the ESP8266 microcontroller (**S2 Appendix**). This module consumes little power (∼ 5V/300mA) and can be powered through a USB port on the board using a generic charger (such as those used for smartphones) or a computer USB. The RGB LEDs on the PCB are controlled via a simple web app developed in HTML/CSS and hosted on the server through which both the intensity and the color of the device’s LEDs are controlled (**Fig 3B and C**). The web page developed is user-friendly and allows, by horizontal slider-type controls, to control the RGB components of the color of the LEDs, each one represented by an integer from 0 and 255, including overall LED intensity (**Fig 3C**). The firmware and software to control this PCB are described in supplementary information (**S2 Appendix**). The filter slider consists of three lee filters, two carefully selected to transmit green (#105) and red (#113) fluorescence emission with minimum background and one that was inappropriate for both wavelengths (#122) [44](**Fig 3D**). The software as well as editable design files (openSCAD) required to reproduce the device are openly available in a GitLab repository [45] as well as a student guide (**S3 Appendix**).

**Figure 3:**
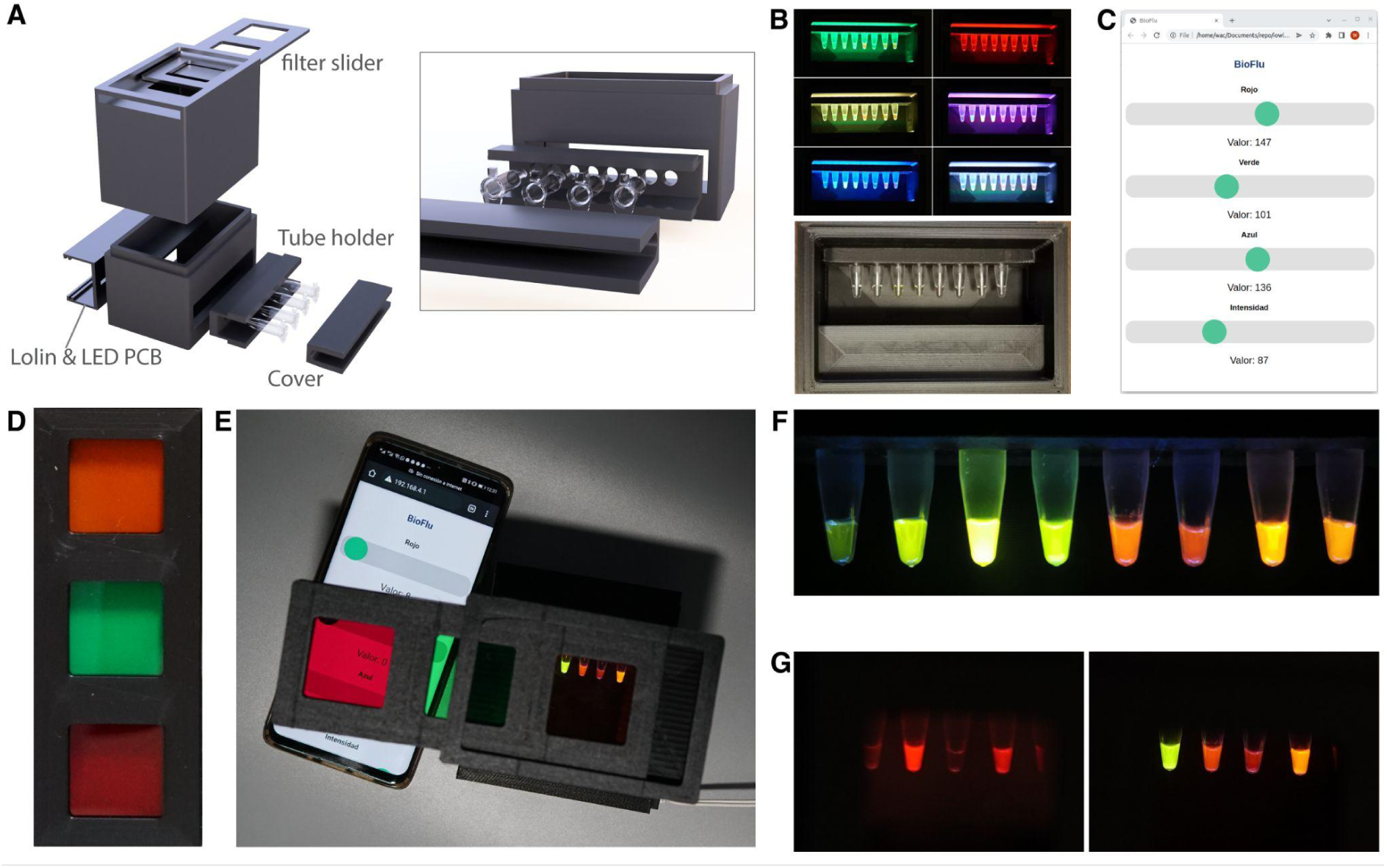
RGBfluor. a) Schematic representation of the 3D printed components of RGBfluor. b) Examples of illumination wavelengths (filter slider removed for clarity). c) User interface to control the LED-based RGB illumination from a mobile phone or external computer. d) 3D printed filter slider with low-cost lee filters. e) Image of the RGBfluor device in use. f) Blue illumination and orange filter setup for different fluorophores, from left to right: 0.01µM FITC, 1µM FITC, 10µM FITC, sfGFP, mScarlet-I, mBeRFP, CyOFP and RRvT. g) Comparison of sfGFP, mScarlet-I, mBeRFP, and CyOFP fluorescence, imaged with green light and red filter (left) and blue light and orange filter (right). Unprocessed raw images; no post-editing applied.

RGBfluor is able to work with a wide range of green-to-red wavelength emitting fluorophores. An example illustrating different fluorescent proteins and dyes under long-pass orange emission filter and blue illumination is shown as an example in **Figure 3E and F**. In this practical, students learn to distinguish between two groups of fluorophores, in the red and green spectra. For instance, students are challenged to take an image with minimum background for all the provided proteins, for which the concepts of long pass filtering and long Stokes shift are applied (**S3 Appendix**). Next, they are asked to selectively image the red fluorescent protein, for which a red filter and green light is preferred to cut off any blue or green light (**Fig 3G**). By the end of this practical, students were expected to gain a comprehensive understanding of fluorescence measurement principles, optimize the selection of filters and illumination conditions for enhanced signal-to-background ratios, and apply these techniques to accurately differentiate between fluorophores in biological samples. This practical was less complex than previous exercises, as it mainly involved the analysis of pre-prepared samples by the students. Overall, all students successfully met the objectives of the activity

### Practical on enzyme kinetics

Our course also consists of a module dedicated to enzyme kinetics, a subject that integrates concepts from different fields such as physical chemistry, biology and mathematics [46]. We developed a hands-on practical on enzyme kinetics using low-cost, locally produced resources. To study the Michaelis-Menten-like kinetics of an enzyme, we use the beta-galactosidase enzyme from *E*. *coli* and the substrate o-nitrophenyl-β-D-galactopyranoside (ONPG), which produces o-nitrophenol, a compound quantifiable by absorbance at 420 nm (**Fig 4**). The β-galactosidase enzyme used in this practical was produced via a cell-free transcription-translation coupled (Tx-Tl) reaction [47], enabling low-cost production as a GMO-free expression system that can be lyophilized and transported at room temperature until rehydration. Users start from freeze-dried reactions, which they rehydrate and aliquot into tubes containing different amounts of ONPG substrate. The reactions are incubated for 12 min at 37°C, and the yellow product is quantified using three methods: image analysis, absorbance measurement using an open-source colorimeter, or a commercial spectrophotometer (**Fig 4A and S4 Appendix**).

**Figure 4:**
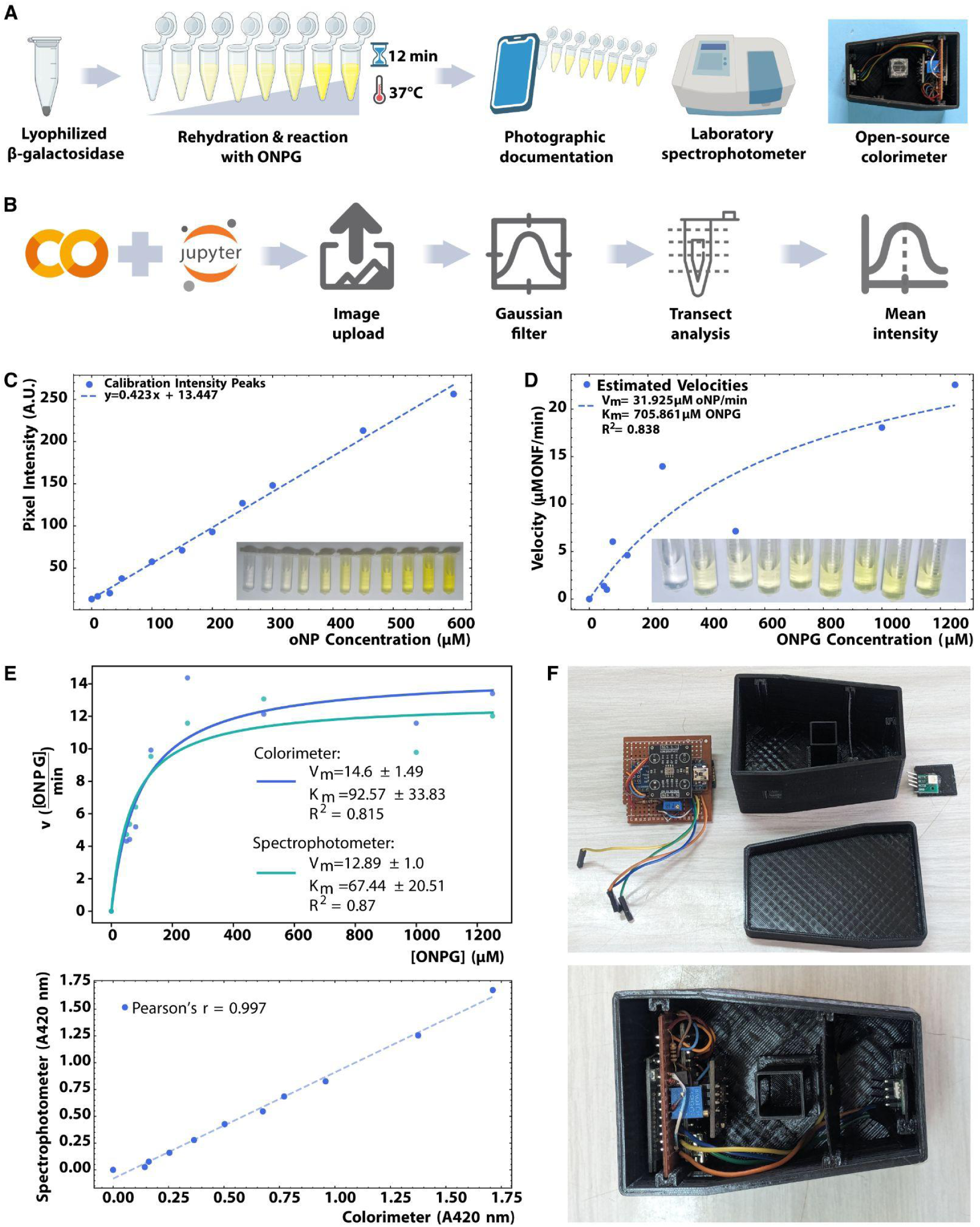
Enzyme kinetic assays. a) Schematic representation of the practical on enzyme kinetics, using a lyophilized and cell-free produced enzyme and two different measurement methods: phone image analysis and absorbance measured on an open source colorimeter, both compared to commercial spectrophotometer. b) Workflow of image processing with Jupyter notebooks in the Google collaborative platform. c) Calibration curve: ONP intensity peaks vs concentration. Inset, image of tubes containing a serial dilution of ONP used to build the calibration curve. d) Estimated velocities vs ONPG concentration Michaelis-Menten curve. Inset, image of tubes used for processing. e) Enzyme kinetic assay with an open source colorimeter in comparison to a commercial spectrophotometer. Top, Michaelis-Menten model fitting. Bottom, correlation analysis between the two approaches. f) Open source colorimeter composed of an arduino nano, bluetooth module, LED, TCS3200, a LED constant current regulator and a 3D printed case.

For the image processing approach, we utilized Google Colaboratory Jupyter notebooks custom-designed for this practical (Enzyme Kinetics: Build a kinetic curve from phone images from [34]) to filter the images and quantify intensities along digital transects across the tubes (**Fig 4B**). Two images were used: one representing a series of o-nitrophenol dilutions to build a calibration curve from the image intensities, and the other depicting tubes containing β-galactosidase and ONPG at different concentrations. Both images were processed to remove the background and noise, facilitating the quantification of intensity. By selecting a transect from each image, we were able to fit a linear regression to the calibration curve and a non-linear Michaelis-Menten function to the saturation curve, yielding parameters within the expected range [48] (**Fig 4C and D**).

For the method based on an open source colorimeter, we used the “reGOSH Colorimeter” [49] (**S5 Fig**). This device integrates a TCS3200 sensor and an RGB LED or, alternatively, a near-UV LED (depending on the application) directly onto its printed circuit board (PCB). Components were chosen based on their local availability and low-cost. The firmware and the software for Windows is from IORodeo [50], whereas the software for GNU/LINUX has been custom-made. Additionally, an Android-compatible GUI was developed. by CoSensores group [51] for its operation from a mobile phone via Bluetooth. Commercially available LEDs generally do not come with their technical specifications. Therefore, in order to verify the dominant wavelength of the LED, we developed a 3D printed adaptor to measure the emission spectrum of LEDs using a commercial spectrometer (**S6 Fig**) [52]. After testing the samples with both the colorimeter and an expensive spectrophotometer, we observed an almost perfect correlation (Pearsońs r=0.997, **Fig 4E**). By quantifying the amount of product per time as a function of the amount of initial substrate, users are able to extract the kinetic parameters of interest (in this case, the relative maximum velocity of the reaction Vmax and the Michaelis constant Km). These parameters are obtained using a custom Google Colaboratory Jupyter notebook we developed to enable interactive data analysis, application of enzyme kinetics concepts, and visualization of enzyme behavior. Detailed documentation is available at “Enzyme Kinetics: Build a kinetic curve from phone images” from [34]. This practice introduced students to enzyme kinetics, focusing on the analysis of Michaelis-Menten reactions using low-cost resources. Students extracted kinetic parameters (Vmax, Km) through image analysis and open-source colorimetry. The simplified setup, using pre-prepared materials, allowed students to focus on data analysis; all groups successfully achieved these objectives.

### Lab-in-a-box: a set of laboratory equipment and reagents for remote practicals

The lockdowns during the COVID-19 pandemic forced us to adapt our practical courses to an online remote teaching format including lab-based activities such as instrument handling, pipetting, and experimental design. Replicating practical experiences at home requires specialized equipment (e.g. microscopes and thermocyclers) and expensive reagents (e.g. enzymes and buffers) that are challenging to deploy [53]. To address this challenge, we created a box that contains and organizes the lab equipment and reagents for practicals, that organizes the equipment and reagents for practicals and is durable enough for transport between homes. In collaboration with the School of Design at our university, we explored cardboard versions of the box with recycling as a design priority. However, the need for strict sterilization procedures forced us to employ plastic boxes that are more appropriate for sanitization based on spraying and wiping. Thus, we developed a lab-in-a-box based on commercial plastic storage boxes. The lab-in-a-box also contained three smaller plastic boxes inside with all the necessary reagents for each student in the group, along with a set of commonly-used tools (e.g. hardware, sanitizer) (**S7 Fig**). The smaller boxes were chosen with the intention of acting as water baths. Since the main goal for the lab-in-a-box was to enable lab experiences with minimal or no need for household goods, a metallic tray was provided to be used as a sanitizable bench and spillover container. A 3D printed light holder was attached inside the box to hold a white light LED tube and facilitate autonomous illumination. We created a total of fifteen boxes and split the class into groups of three students (one lab-in-a-box per group). Thus, the lab-in-a-box provided a set of tools for common use for the group, while the smaller boxes provided reagents for each student. The rationale behind this organization was to avoid cross-contaminations among students and consider the three experiments separately as technical replicates in the practicals.

In 2020, the distribution of the lab-in-a-box kits was organized to reach each of the 45 students’ homes (3 students per group, 15 groups in total). In 2021, as health safety measures allowed students to return to the laboratory but at a limited capacity, the experiments were conducted in person with social distancing protocols (3 students per group, 16 groups in total). The box setup used for in-lab activities mirrored that of the home-based practicals, with each student in the group conducting the experiment individually over a three-day period, spaced one week apart. Thus, during 2020 the box was sent from the university to the home of the first student in the group, and then passed on to the second and third student before being shipped back to the university for thorough sanitization and refilling. This project reached a geographically diverse group of 45 students across Chile, including both urban and rural areas, and 48 students in teaching laboratories in 2021. Approximately 75% of the participants were from various locations within the Santiago Metropolitan Region. The northernmost delivered box reached Arica, approximately 2,031 km from the distribution point, while the southernmost delivery reached Puerto Montt, approximately 1,033 km away from the university. Local courier services were used to deliver the kits, with adjustments made for each region. While urban students generally received their kits on time, minor delays were encountered in rural areas due to logistical challenges. These were anticipated and managed through prearranged communication with the students, ensuring all could complete their experiments within the required timeframe.

Each student followed detailed video and written instructions for disassembling, sanitizing and reassembling the box before passing it to the next home [54] (**S5 Appendix**). Since most students had no prior experience in molecular biology or pipetting, they were first instructed to complete a micro-pipetting exercise using food coloring (**S6 Appendix**) before starting the practicals.

### The use of OER in high school education

These open educational resources (OERs) could offer a cost-effective approach to enhancing biology learning in high schools. To assess this potential, we organized three workshops for high school teachers from different institutions across Chile. The workshops consisted in hands-on activities with these resources. To evaluate the impact of these resources, we made a questionnaire with response scores on a 1–5 Likert-type scale, ranging from “strongly disagree” (1) to “strongly agree” (5), with following statements:

(Q1) The use of open and low-cost devices helps to better understand both the technique and the underlying concepts.

(Q2) I will likely use some of these resources in the future.

(Q3) The use of open reagents and hardware broadened my perspective on the potential applications of the techniques in my own teaching.

(Q4) I prefer hands-on activities with individual low-cost devices (one per person) rather than demonstrations using a single piece of equipment for everyone.

(Q5) The provided materials were sufficient to understand and carry out the experiments.

(Q6) The use of open and open-source technology facilitates the teaching of molecular biology topics covered in this workshop.

The results from this questionnaire are summarized in **Figure 5**. Overall, the responses were positive towards the use of these OER for teaching in high school settings, with Q2, Q3 and Q6 with 100% responses for “strongly agree” with the statements and the rest of statements being evaluated with average scores above 4.0 (“agree” to “strongly agree”). Q1, “The use of open and low-cost devices helps to better understand both the technique and the underlying concepts” scored 4,33 (s.d.=1,63), Q4, “I prefer hands-on activities with individual low-cost devices (one per person) rather than demonstrations using a single piece of equipment for everyone”, scored 4,33 (s.d.=0,81) and Q5, “The provided materials were sufficient to understand and carry out the experiments”, scored 4,66 (s.d.=0,81).

**Figure 5:**
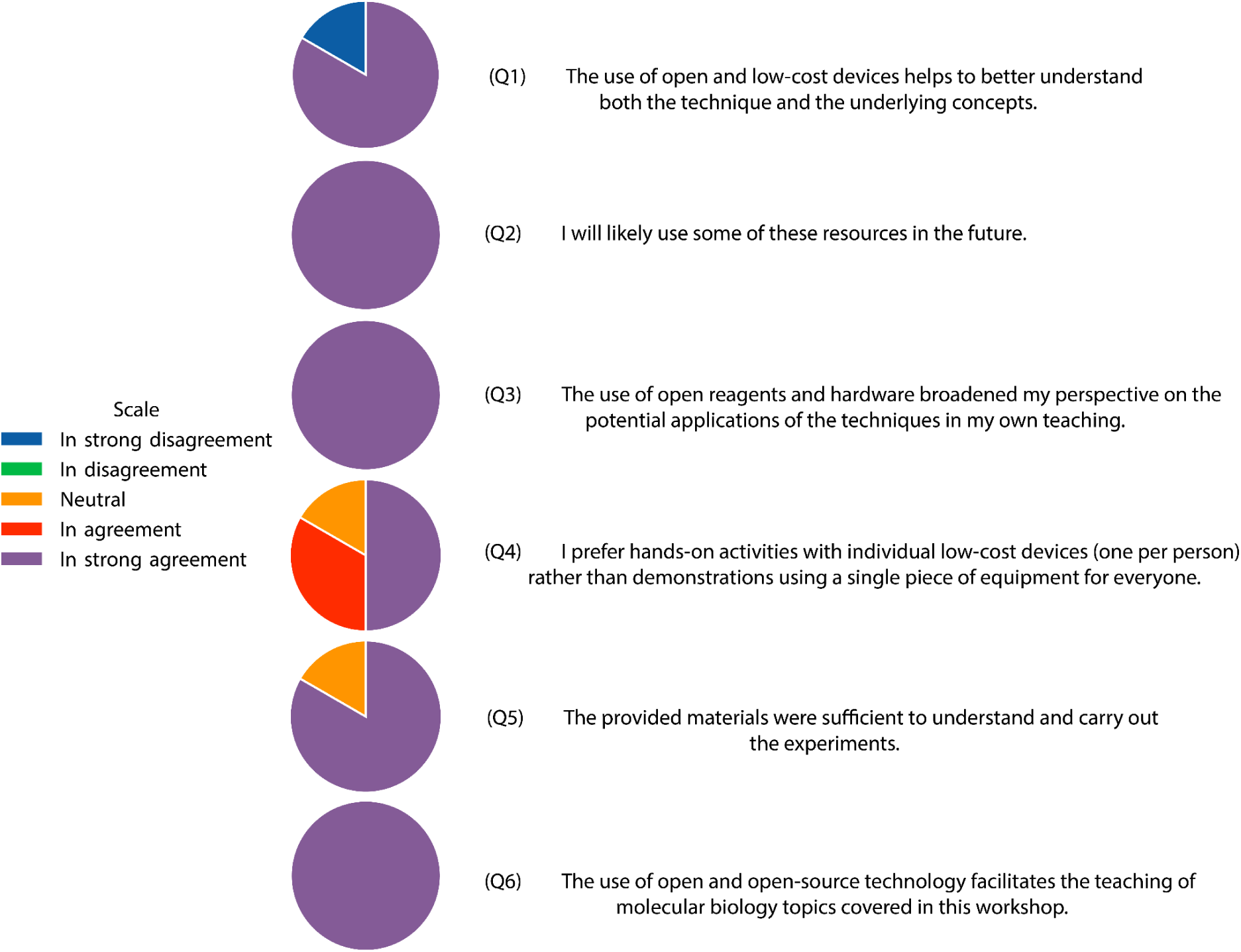
Results from the teachers’ evaluation of OER. A Likert-type scale questionnaire composed of 6 questions was provided to teachers and answered anonymously (n=6). Q1: 4,33 (s.d.=1,63), Q2: 5,00 (s.d.=0), Q3: 5,00 (s.d.=0), Q4: 4,33 (s.d.=0,81), Q5: 4,66 (s.d.=0,81) and Q6: 5,00 (s.d.=0).

In general, these results indicate that these resources were highly valued, and their use in high school teaching is very likely (Q2). Notably, there was strong agreement on the extent to which these open educational resources (OER) broaden teachers’ perspectives on the potential applications of the techniques (Q3), as well as how the use of open and open-source technology facilitates the teaching of molecular biology topics (Q6). The statement with less agreement scores 4,33 related to how the use of open and low-cost devices helps to better understand both the technique and the underlying concepts (Q3), and the preference for hands-on activities with individual low-cost devices (one per person) rather than demonstrations using a single piece for the whole course (Q4). These lower scores, although still in the “agree” level (>4,00), may reflect that further work is still needed to make these resources more intuitive and simpler to use in order to replace traditional tools.

## Discussion

Fostering access to OER contributes directly to the Sustainable Development Goal 4 (SDG4), focused on ensuring education and learning opportunities for all [55]. In the same way community-driven efforts on open source hardware have proven effective for emergency response during the COVID-19 pandemic [55,56], this approach can be employed to achieve more equitable, inclusive and effective access to educational resources. Although the adoption of open tools is growing in academia and is being proposed as an important component of the UNESCO Open Science Recommendation [57], its introduction in the curricula of both university and high school education remains very scarce in our region. The goal of using open-source tools in the OER described here is twofold: to reduce costs and foster autonomy for more sustainable educational models, and to engage students with open-source technology, encouraging discussions on how this approach can drive a paradigm shift toward a more collaborative scientific ecosystem and open innovation models in the future [58,59]. The local production of both hardware and reagents significantly lowers the costs of practical teaching. For instance, the homemade production of cell free reactions, RT-PCR and LAMP results in cost reductions of one to two orders of magnitude for our institution [20,60]. Further reductions could be achieved in the future, for example, with the use of cellular reagents [61] to replace enzyme purification protocols. The cost reductions associated with these OERs are also enabling their employment in other courses that previously lacked practical sessions due to financial constraints. For instance, they enabled the creation of new hands-on modules in two other courses on Synthetic Biology and Open Hardware prototyping. We have also explored their potential use in high schools through training workshops for high school teachers in Patagonia who positively assessed their implementations (Fig. 5). There was strong agreement on how these OER broaden teachers’ perspectives and facilitate the teaching of molecular biology topics.

Complying with Open Science [57], OER [62] and open hardware recommendations [63], we provide detailed instructions, schematics, bill of materials and editable design files in openly accessible online repositories with version control (git). These resources are released under open licenses that grant students and teachers the freedom to replicate these resources and build upon this work. Protocols for the production of buffers and enzymes were also deposited in an openly accessible repository, protocols.io. The use of free software in the form of shareable notebooks introduces students not only to a growing trend in data analysis and programming [64], but also to the relevance of good practices in software documentation to enable reproducibility of their results. Reproducibility is a core pillar of Open Science [57]; and open access computational tools such as Jupyter notebooks are essential, as they allow others to reproduce results reliably [65]. Jupyter Notebooks foster reproducibility by integrating executable code, documentation text and analysis output in a single place that can be read and re-run by other users. We have used Jupyter notebooks in classrooms both as standalone simulation environments and as platforms for data interaction and analysis [41]. As these notebooks become readily available upon opening and do not require time-demanding installations of software libraries and dependencies, their use leads to more fluid class experiences [66].

Another key goal of this work was to deepen students’ understanding of the underlying mechanisms behind the instruments and scientific principles used. Here, using open-source hardware (OSH) as individual mini-labs provides a significantly improved learning experience compared to traditional setups, such as 50 students crowded around a single’black box’ microscope to learn the principles of fluorescence. We are currently working to expand this approach by incorporating devices for DNA assembly [67], fluorescence microscopy [41], optogenetics [68][69] and integrated incubation and real time reading from reactions [70]. We also aim to implement the Airflow microreactor for isothermal DNA amplification [71], as well as ultra low-cost devices for isothermal incubation [72].

An important test for these OER was to address the challenge of remote teaching during the pandemic lockdowns. The COVID-19 pandemic forced many universities around the world to close their facilities and abruptly adopt remote teaching, leading to an urgent need for online teaching resources. In response to this challenge, educators across different fields developed novel teaching resources. While remote teaching has long been part of educational strategies [73], the lockdowns to mitigate SARS-CoV-2 spread led to an unprecedented growth of at-home teaching resources. For instance, cloud-based tutorials [66] and virtual interactive classroom [74] have been developed for bioinformatics courses in life sciences. However, practical courses that primarily rely on laboratory experimentation faced a more complex scenario as hands-on experiences are difficult to replace by online tools [73]. Despite the lack of preparedness for such an unprecedented challenge, pre-pandemic developments such as online programming notebooks [75], low-cost biological resources for advanced practicals in molecular biology [24,31,76–79], and a wide range of open source hardware devices for scientific research [50,80,81] had paved the way for the implementation of remote teaching in molecular biology. In this context, ‘remote laboratories’ refer not to centralized facilities operated remotely [82], but rather to the deployment of instruments and reagents directly to students’ homes. Efforts to democratize biotechnology, led mainly by biomakers, open community labs, do-it-yourself biology practitioners and advocates for open biotechnology, have provided resources and documentation for hands-on practical workshops, and enabled low-cost experimental research outside university laboratories [8,9,83–86]. These initiatives provide not only valuable guidelines for frugal biotechnology development but also stringent biosafety and biosecurity regulations for their deployment [87]. These efforts have been essential for the development of our lab-in-a-box in a couple of months, with reaction success rates ranging between 73% and 100% across different groups. These positive results demonstrate the robustness of these resources, given that students had no prior pipetting training, the samples were exposed to high temperatures (i.e., transparent box acting as a greenhouse upon direct sunlight exposure during transport) and there were no second chances if any device or reagent failed upon delivery.

The use of open source hardware has also triggered new collaborations with emerging open hardware companies, such as IORodeo, GaudiLabs and IOWlabs, as well as academic initiatives such as GMO detective and with members of Gathering for Open Science Hardware (GOSH). The collaboration with IOWlabs has led to the co-design of devices that are scaled and sold to the university while they remain free to be studied, adapted and commercialized by third parties. The collaboration with GMO detective, for instance, has led to a contribution to their design, later improved by members of the Hackteria network. This open interaction between academic laboratories and the private sector has given rise to a highly efficient and fluid collaborative model that has been key for a rapid response to these lockdowns (i.e. approximately 4 months to obtain all the resources for the whole course). We aim to work towards a future scenario in which researchers create, share and build their own instruments in collaboration with commercial initiatives that can scale up production based on their own developments or upon other designs. This change in innovation approaches would require nurturing future generations by addressing the topic of OSH in the curriculum [59]. Integrating OER into curricula represents a foundational step toward embedding open source principles in scientific education and the development of collaborative research ecosystems. These could complement other efforts such as online teaching resources [79][88][89], and educational problem-solving initiatives [90][76].

## Materials and methods

### Course Description and Structure

The course was *Biochemistry Laboratory II: Molecular Genetics*, aimed at third-year biochemistry students in the Department of Biological Sciences at Pontificia Universidad Católica de Chile. The data presented here correspond to two years, 2020 and 2021. In 2020, due to health restrictions, the course was conducted entirely online, with materials sent to the students’ homes. In 2021, the course returned to an in-person format with social distancing protocols in place. Students were organized into groups of three, with each student conducting the practicals independently. The course spanned one academic semester, and each practical session was designed to take no more than three hours to complete, whether conducted online or in person. After all members of a group completed their experiments, they discussed their results and prepared a written report, followed by an oral presentation where the results were openly discussed with the entire class. These reports and presentations formed the basis for the students’ evaluations. The course had a total of 45 students in 2020 and 48 students in 2021. Each year, four teaching assistants (TAs) were assigned to prepare all the materials and provide support, either remotely (via Zoom) or in person, ensuring that the experiments worked properly and addressing any questions the students had. All students were provided with detailed documents (**S5 Appendix**) and given safety briefings on biosafety measures and social distancing protocols to maintain a safe working environment.

## Reagents preparation

### RT-PCR for SARS-CoV-2 Detection

The positive RNA control for the RT-PCR assay was synthesized using a standardized protocol based on the procedures detailed in [20]. Initially, the pT7_N_amplicon template, derived from the SARS-CoV-2 N gene positive control vector (IDT, Cat # 10006625), was amplified using the *NT_Fw* and *T7_Nter_Rv* primers. The resultant PCR products were subsequently subjected to purification through agarose gel electrophoresis employing the Wizard Sv Gel Clean-up system (Promega). Following purification, the products were used to generate RNA transcripts using the HiScribe T7 In Vitro Transcription Kit (NEB, E2040S) according to the manufacturer’s instructions. Post-transcription, the DNA template was removed by treatment with DNase I (NEB), followed by purification of the RNA transcripts using the RNeasy Kit (Qiagen) as per the supplied protocol. This procedure yielded viral N RNA suitable for use as a positive control in RT-PCR assays (∼50 μL at 1,000 ng/μL). The construction of vectors encoding the enzymes used in this study has been previously described [20]. Additionally, all proteins expressed in this work were purified following detailed protocols, which have been optimized and documented. These protocols are publicly accessible on the protocols.io platform [26,37]. RT-PCR was conducted with N1 and N2 primers for the SARS-CoV-2 N gene (**Table I**), as recommended by the CDC’s Research Use Only primers list [91]. Reactions were performed using the previously described BEARmix reaction buffer [19]. Each RT-PCR reaction mix was prepared in a total volume of 20 μL, consisting of the following components: 1 μL Pfu-Sso7d (0.3 mg/mL), 0.125 μL M-MLV RT (0.02 mg/mL), 2 μL DTT (100 mM), 2 μL each of forward and reverse primers (3 μM, N1 or N2 as appropriate), 4 μL 5X BEARmix Buffer, 2 μL dNTPs (4 mM), 3.875 μL nuclease-free water, and 5 μL RNA sample. To each tube, 10 μL of mineral oil was added without mixing to prevent evaporation of the reaction mixture. The reactions were then incubated in the PocketPCR thermocycler (https://gaudi.ch/PocketPCR/) following a program set at 50°C for 15 m, followed by 30 cycles of 98°C for 10 s, 56°C for 20 s, and 72°C for 30 s, with a final extension step at 72°C for 5 m. To visualize DNA bands, a 2% agarose gel stained with SYBR™ Safe DNA Gel Stain (Thermo Fisher Scientific) was employed, and loaded alongside a 50 bp DNA Ladder. Electrophoresis was conducted in a 1X TAE buffer using the BlueGel™ electrophoresis system for 30 min [91].

**Table I.**
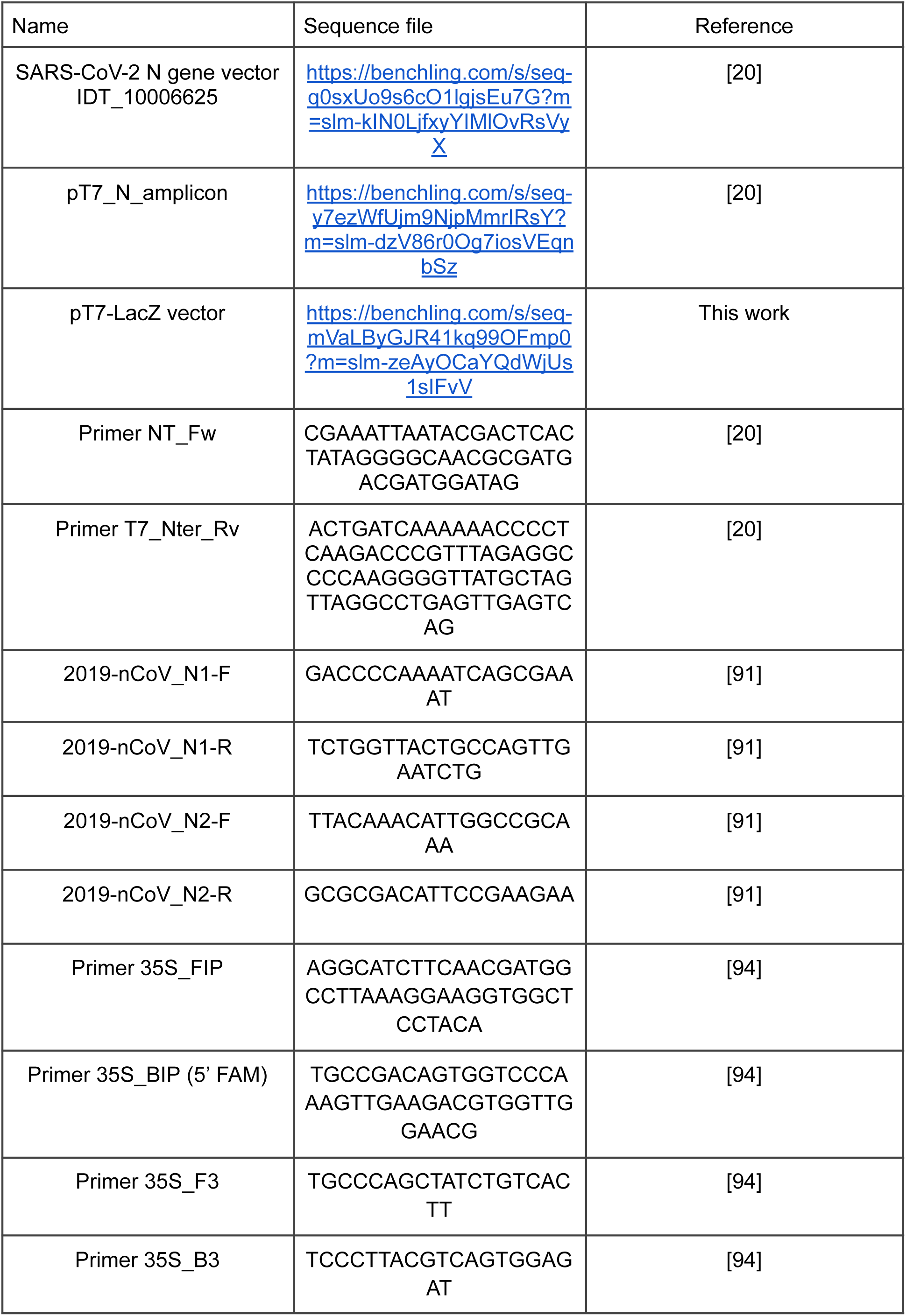
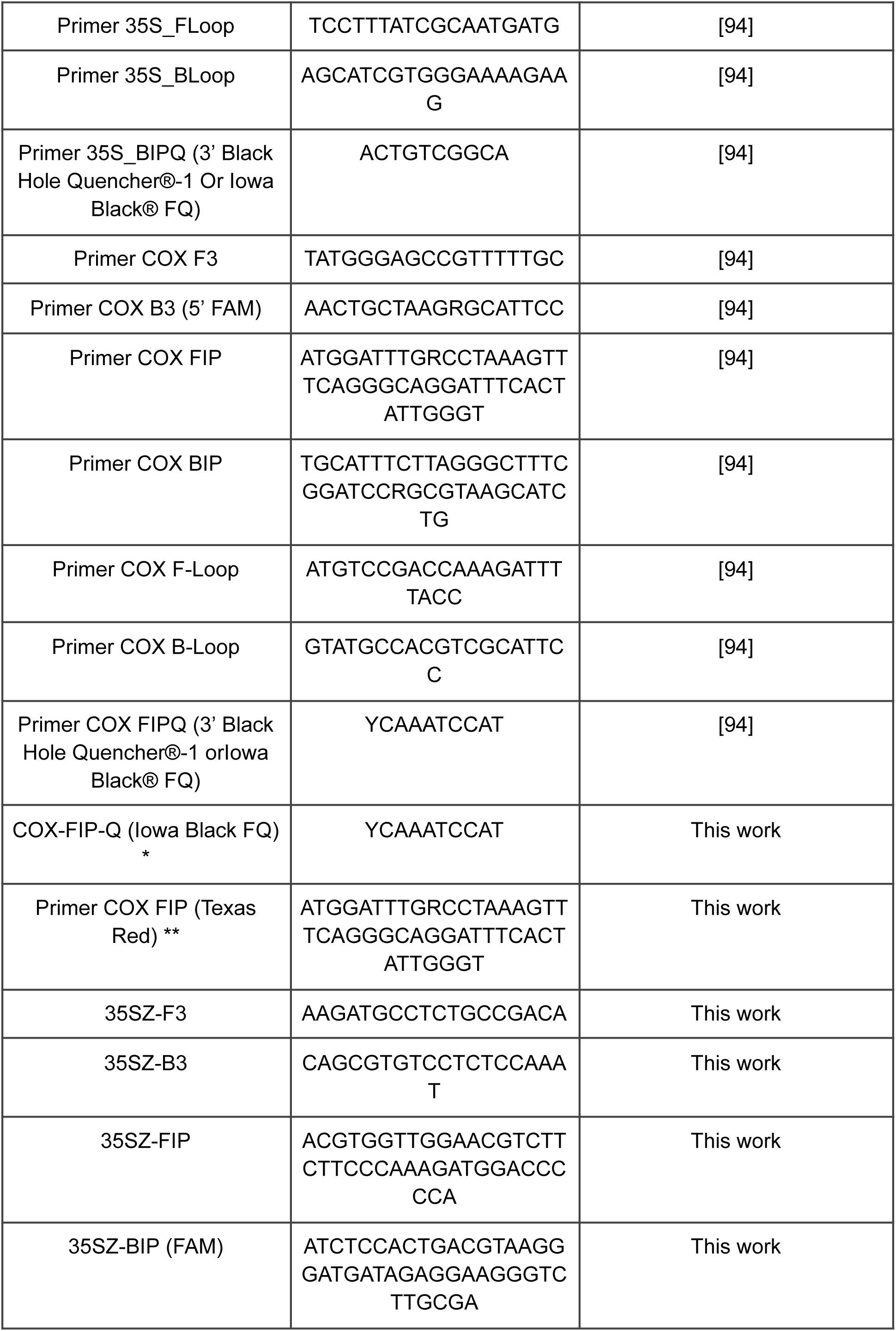
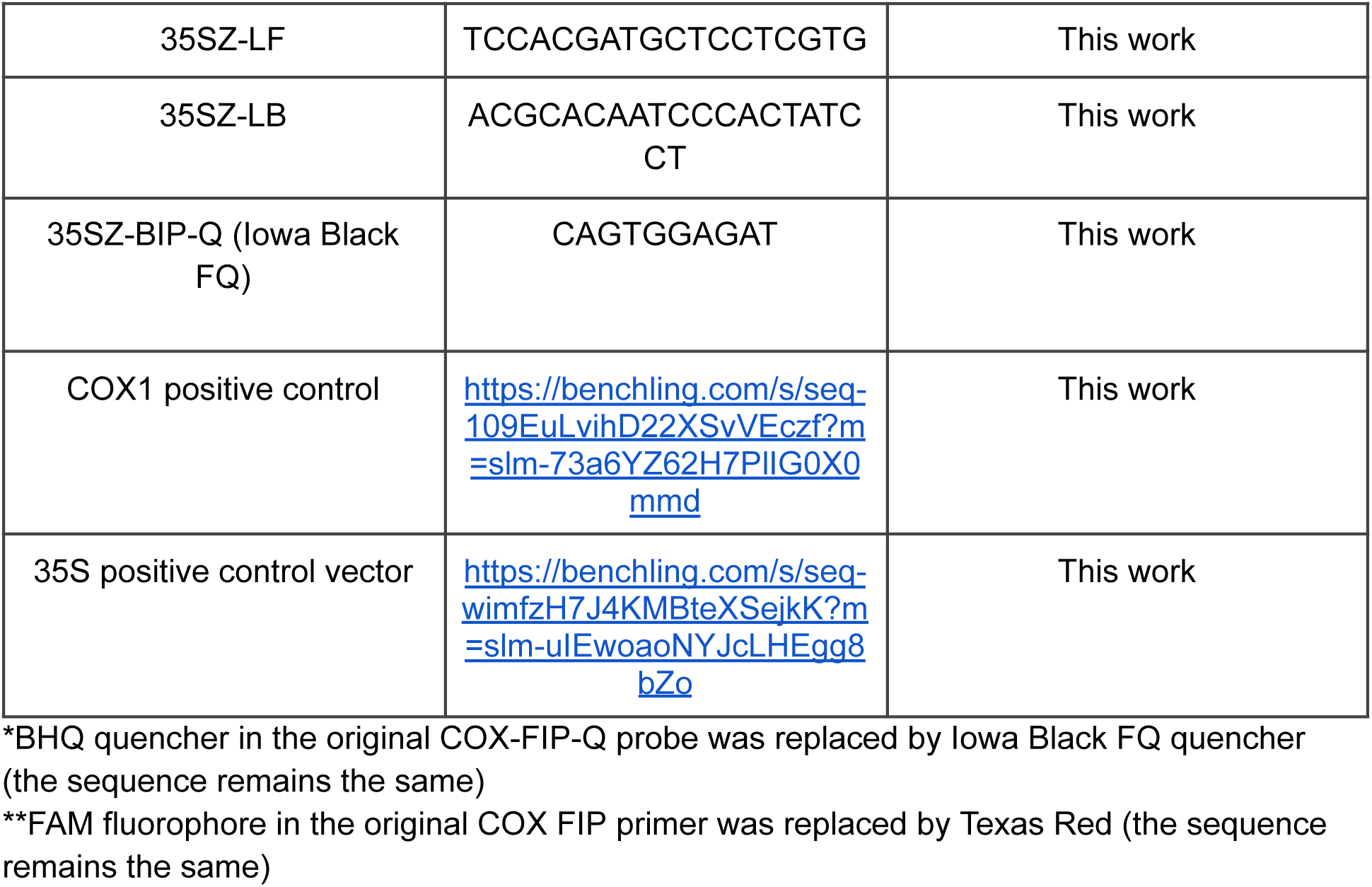
DNA sequences.

### DNA Detection via LAMP Assays

Fresh LAMP reactions were prepared following a previously described LAMP protocol but adapted for QUASR detection [21]. In short, 10 µl homemade reactions contained of 1X Isothermal Buffer (20 mM Tris-HCl pH 8.8, 10 mM (NH4)2SO4, 50 mM KCl, 2 mM MgSO4, 0.1% Tween 20), 6 mM MgSO4, 1.4 mM of dNTPs, 0.16 mg/mL of locally-produced BstLF polymerase and a LAMP-QUASR 35S primer mix for the detection of the 35S promoter (1.6 µM FIP and BIP-FAM, 0.8 µM LF and LB, 0.2 µM F3 and B3, and 2.4 µM BIPc-Q; **Table I**). Reactions were incubated at 65 C for 35 minutes and then cooled down at 15 C for 15 minutes before being observed under a blue LED light using the acrylic or 3D-printed GMO Detective [39]. Nuclease-free water was used as a negative control and around 20000 copies of the 35S positive control.

For the lyophilization of the homemade GMO Detective reactions, we followed the protocol described online in [38]. The reaction mix was prepared with 0.16 mg/mL *BstLF* (stored without glycerol at-80°C), 1.4 mM dNTPs, a 10X mix of LAMP 35S-Z primers and trehalose at 5% v/v. Each tube with 8 μl of this mix was sealed with parafilm and pre-frozen at-80°C for 10-20 minutes. These steps are important to avoid the content spilling over in the freeze-drying process due to abrupt pressure changes, and also to ensure a solid state of the mix that favors correct sublimation. The parafilm of each tube was perforated (with a sterile syringe) and the preparations were placed in a FreeZone 2.5L dessert freeze dryer (LabConco) at-84 °C and 0.04 mBar for 2-3 hours, gradually increasing the temperature during the last 30 minutes. The tubes were closed and sealed in a vacuum in embossed plastic bags with a silica gel bag and an oxygen absorber. A pulse of argon gas was delivered inside to clear the air from the atmosphere in contact with the freeze-dried material. The bags were deposited in aluminum bags that were also sealed in a vacuum, allowing the preparations to be stored in the dark and at room temperature or 4°C for a period of 7 days to 2 months. Subsequently, each lyophilized mixture was rehydrated with DNA and a mixture containing 6 mM MgSO4 and 1X Isothermal Buffer at a final volume of 10 μl per reaction. The reactions were incubated and their fluorescence was observed in a similar way to the fresh reactions. BstLF enzyme was expressed and purified following the protocol described in protocols.io [37], and diluted to a working concentration of 0.2 mg/mL. A fraction of the enzyme was stored in a buffer with glycerol at-20°C until use in LAMP reactions, while another fraction was stored at-80°C in a buffer without glycerol for use in a master mix of freeze-dried LAMP reactions.

### Cell free reactions

Cell-free expression of the beta-galactosidase enzyme was produced incubating plasmid DNA (pT7-LacZ vector, 10nM) with 33.33% of cell-lysate from a BL21 DE3 strain deficient in the LacZ gene (https://www.addgene.org/99247/) and reaction buffer containing phosphoenol pyruvate as energy source [60]. Freeze-dried steps, when needed, were performed using a-80C benchtop lyophilizer following a protocol detailed in [92] for LacZ.

The colorimetric substrate for quantification of the β-galactosidase activity, o-nitrophenyl-β-D-galactopyranoside was purchased from Sigma-Aldrich (cat N1127) as well as the product o-nitrophenol (cat N19702). Both were diluted to 10 mM o-nitrophenol and 3 mg/ml ONPGal in saline buffer (PBS,137 mM NaCl, 10 mM phosphate, 2.7 mM KCl, pH 7.4) as stock solutions (**S4 Appendix**).

Positive controls for COX1 and 35S were produced by commercial DNA synthesis and cloning into pL0R uLoop vectors [93] (**S1 Appendix**).

### Hardware design and fabrication

All 3D printed parts were designed using the open-source software openSCAD, exported as STLs and printed in Ender 3 or Prusa MK6 3D printers. The openSCAD editable files of GMO device and RGBfluor are available in the git repositories [39] and [45], respectively.

Printed circuit boards were designed using the open-source software KiCAD 5. PCBs were manufactured in the JLC company and the components were acquired through the LCSC company.

### Glossary

OSH: “any piece of hardware used for scientific investigations that can be obtained, assembled, used, studied, modified, shared, and sold by anyone. It includes standard lab equipment as well as auxiliary materials, such as sensors, biological reagents, analog and digital electronic components.” [58]

OER: “Open Educational Resources (OER) are learning, teaching and research materials in any format and medium that reside in the public domain or are under copyright that have been released under an open license, that permit no-cost access, re-use, re-purpose, adaptation and redistribution by others.” [95]

## Acknowledgements

We thank Fernando Guzmán-Chávez and Jim Haseloff for sharing plasmids encoding RRvT. Valerie Decap, Clara Espinola, Javier Alejandro Contreras Castro, and Paulina I. Merino Sepúlveda for the support on teaching labs and material preparation.

We also acknowledge the gracious help of Amparo Nuñez during the protein purification processes in the final stages of this work. We thank members of the gLAMP consortium, ReClone forum (https://reclone.org/), Open Bioeconomy Lab (https://openbioeconomy.org/) and the JOGL OpenCOVID community (https://app.jogl.io/program/opencovid19) for the advice and for sharing relevant information regarding diagnostic reactions for SARS-CoV-2. We also thank the FreeGenes project, Alex Brown, Ali Bektas, Jose Gomez-Marquez, Jim Haseloff, for valuable conversations and advice. Figures were created with BioRender.com. The PCR tube 3D model was created from OpenSCAD code at https://github.com/Biotinkering/PCR-Tube

## Author Contributions

FF, JM, AC, designed research;

FF, JM, GA, AL, CSR supervised research;

AC, AA, VZ, APA, WA, DG, JA, FQ, IN, FN, TM, VF, MB, SV, MR, GA, FC, PC FF performed research;

SA, SR, designed the lab-in-a-box

RO formulation of questionnaire and text editing. AC, AA, APA, FN, and FF analyzed data;

AC, AA, APA, FN, and FF wrote the manuscript. All authors reviewed the manuscript.

RO Ruby Olivares

AC Ariel Cerda

AA Alejandro Aravena

VZ Valentina Zapata

APA Anibal Arce

WA Wladimir Araya

DG Domingo Gallardo

JA Javiera Aviles

FQ Fran Quero

IN Isaac Nuñez

FN Felipe Navarro

TM Tamara Matute

VF Valentina Ferrando

MB Marta Blanco

SV Sebastian Velozo

SR Sebastian Rodriguez

SA Sebastian Aguilera

JM Jennifer Molloy

GA Guy Aidelberg

AL Ariel Lindner

FC: Fernando Castro

PC Pablo Cremades

CRS Cesar Ramirez-Sarmiento

FF Fernan Federici

## Data Availability

All data generated or analyzed during this study is included in this published article and its Supplementary Information files. Additionally, all relevant data has been deposited in a Zenodo repository (https://doi.org/10.5281/zenodo.13623939), organized as follows:

- Raw Figures, 1-4: Original.TIF, JPEG,.PDF and.PNG images used for Figures 1 through 4. In Figures where numerical and/or image data was used, raw datasets in Excel (.XLSX) format and Jupyter Notebooks (.ipynb) are available for reproducing plots.

- Jupyter Notebooks: Original Jupyter Notebooks used. Both Original.ipynb and Google Colaboratory file versions are available.

- Plasmid Sequences: Original and annotated plasmid sequences in GeneBank format (.gb) for the 35S positive control vector, COX1 positive control, SARS-CoV-2 N Gene, pT7-LacZ and pT7_N_Amplicon.

- Protocols: PDF files describing the protocols for Cell-Free Extract expression and lyophilization of beta-galactosidase, Lyophilization and manufacturing of Corona Detective Assay, LAMP reaction protocols, Recombinant expression and purification of M-MLV, Bst-LF, Pfu Sso7d and Mashup.

Correspondence and requests for any materials or additional information should be addressed to the corresponding author upon reasonable request.

**S1 Appendix: Homemade GMO Detective.** Detailed modifications to the original GMO Detective protocol, including custom primer sets, QUASR probes, and comparison of LAMP reactions using different fluorescence labels.

**S2 Appendix: RGBfluor instructions.** Guide to the design, assembly, and operation of the RGBfluor PCB, including detailed schematics, bill of materials, and firmware setup for controlling RGB LEDs via Wi-Fi.

**S3 Appendix: RGBfluor student guide.** A practical guide for students using the RGBfluor device in fluorescence experiments, covering learning objectives, setup instructions, and tasks for detecting fluorescent proteins. Includes links for fluorescent protein profiles, emission filters, and design files.

**S4 Appendix: Enzyme kinetics student guide.** Instructions for a hands-on experiment to characterize the kinetic parameters of β-galactosidase from *E. coli*.

**S5 Appendix: Student guidelines for using and sanitizing the lab-in-a-box**. A guide for safely receiving, using, and handling the lab-in-a-box materials, including sterilization procedures, biosecurity standards, and instructions for both LAMP and RT-PCR practicals.

**S6 Appendix: Student instructions for micropipetting**. A guide on how to correctly use micropipettes, covering their parts, types, and proper techniques for accurate liquid handling. This appendix includes practical exercises for the subsequent experimental activities.

## Supplementary information

**Figure S1:**
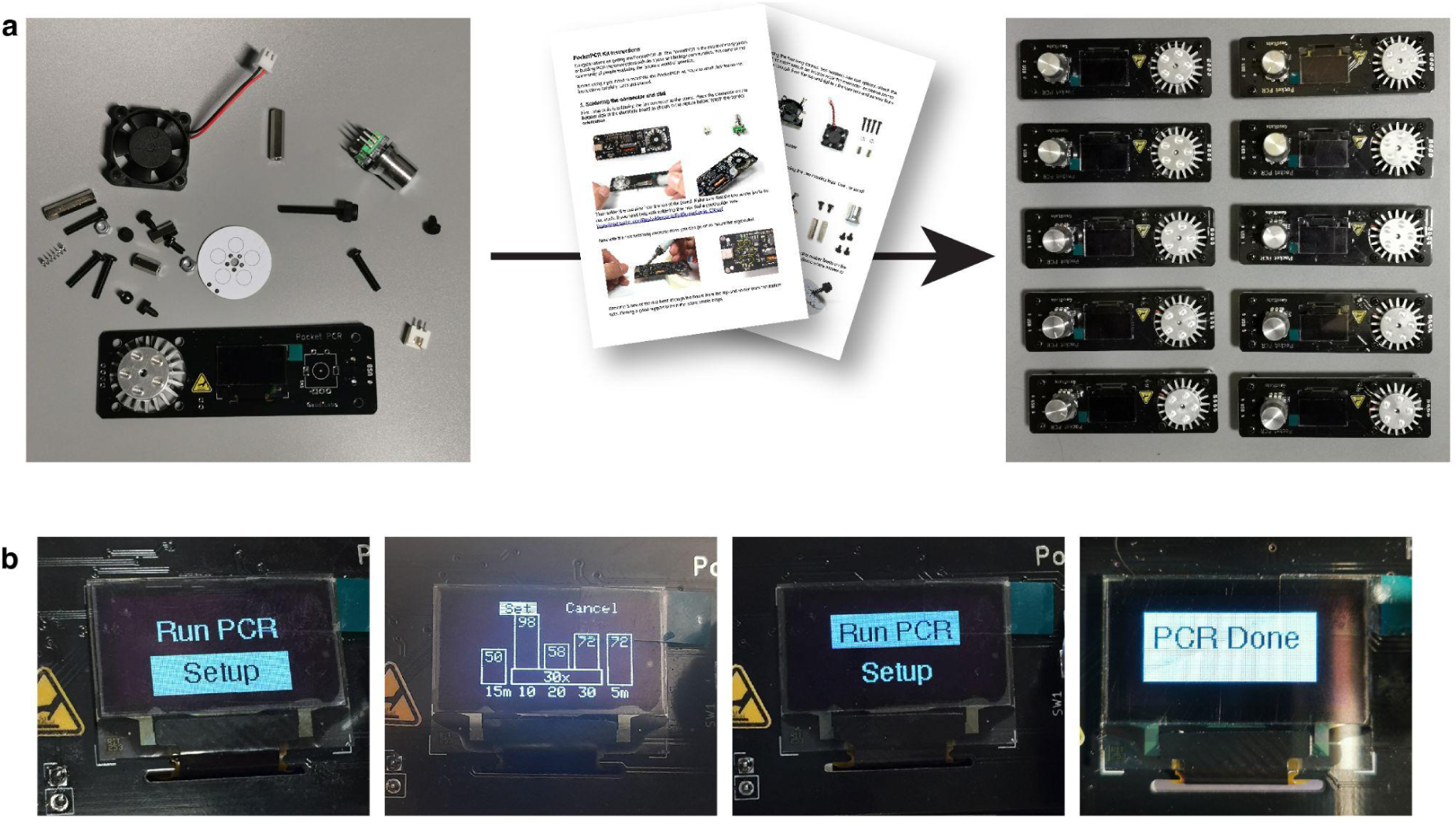
**PocketPCR assembly and operation**. a) PocketPCR is sold as a kit to be assembled with minimum soldering (it can also be obtained fully assembled). Clear instructions are provided by GaudiLabs. b) Sequential steps showing thermocycling setup. The main knob is rotated to choose values and pushed to confirm selections.

**Figure S2:**
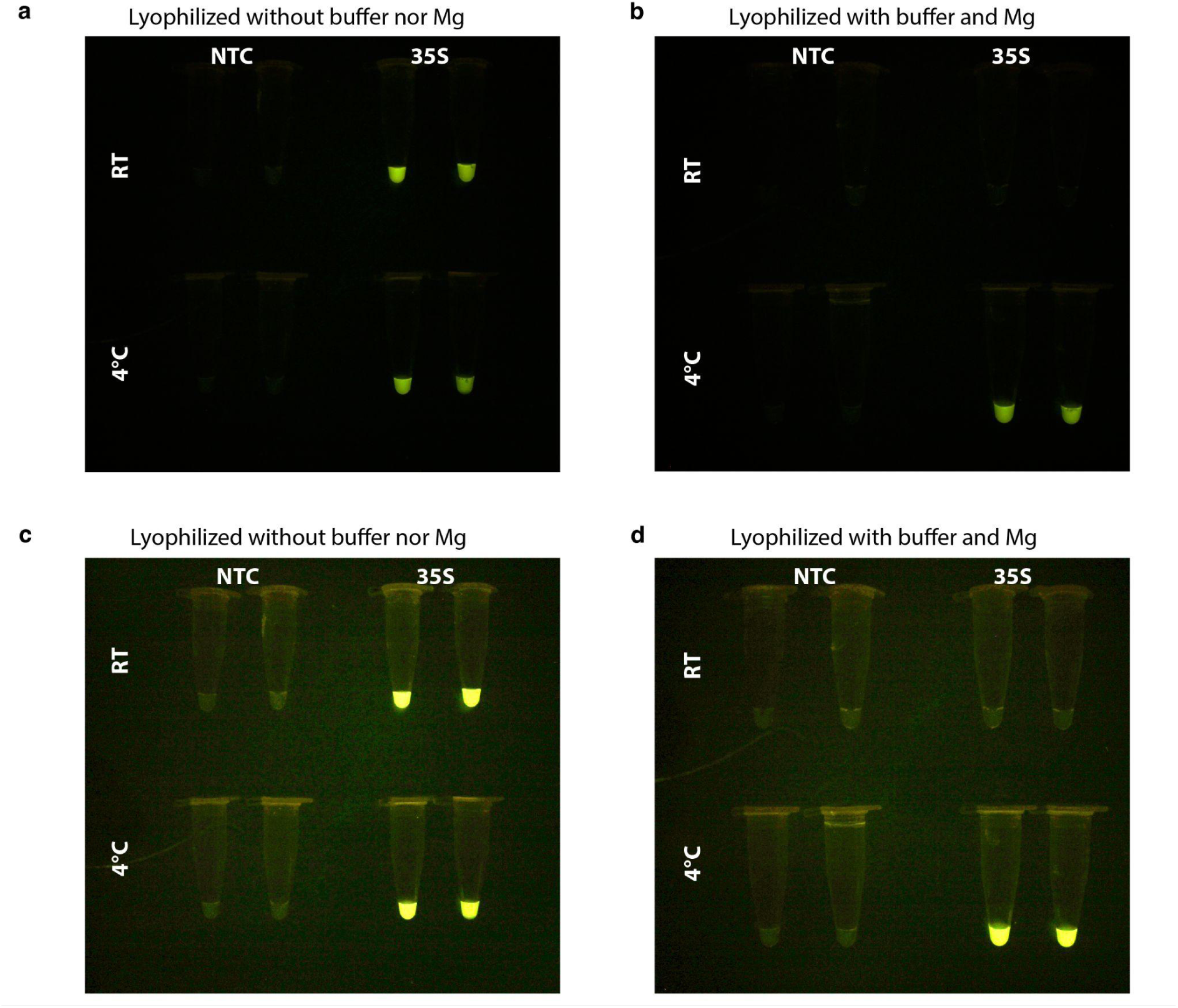
Lyophilization without magnesium nor reaction buffer provides longer storage capabilities. a) and c) Reactions lyophilized without buffer nor magnesium, rehydrated after two-month storage at room temperature (top) or 4°C (bottom). b) and d) Reactions lyophilized with buffer and magnesium, rehydrated after two-month storage at room temperature (top) or 4°C (bottom). Panel c and d are images with overexposure to show the negative tubes.

**Figure S3:**
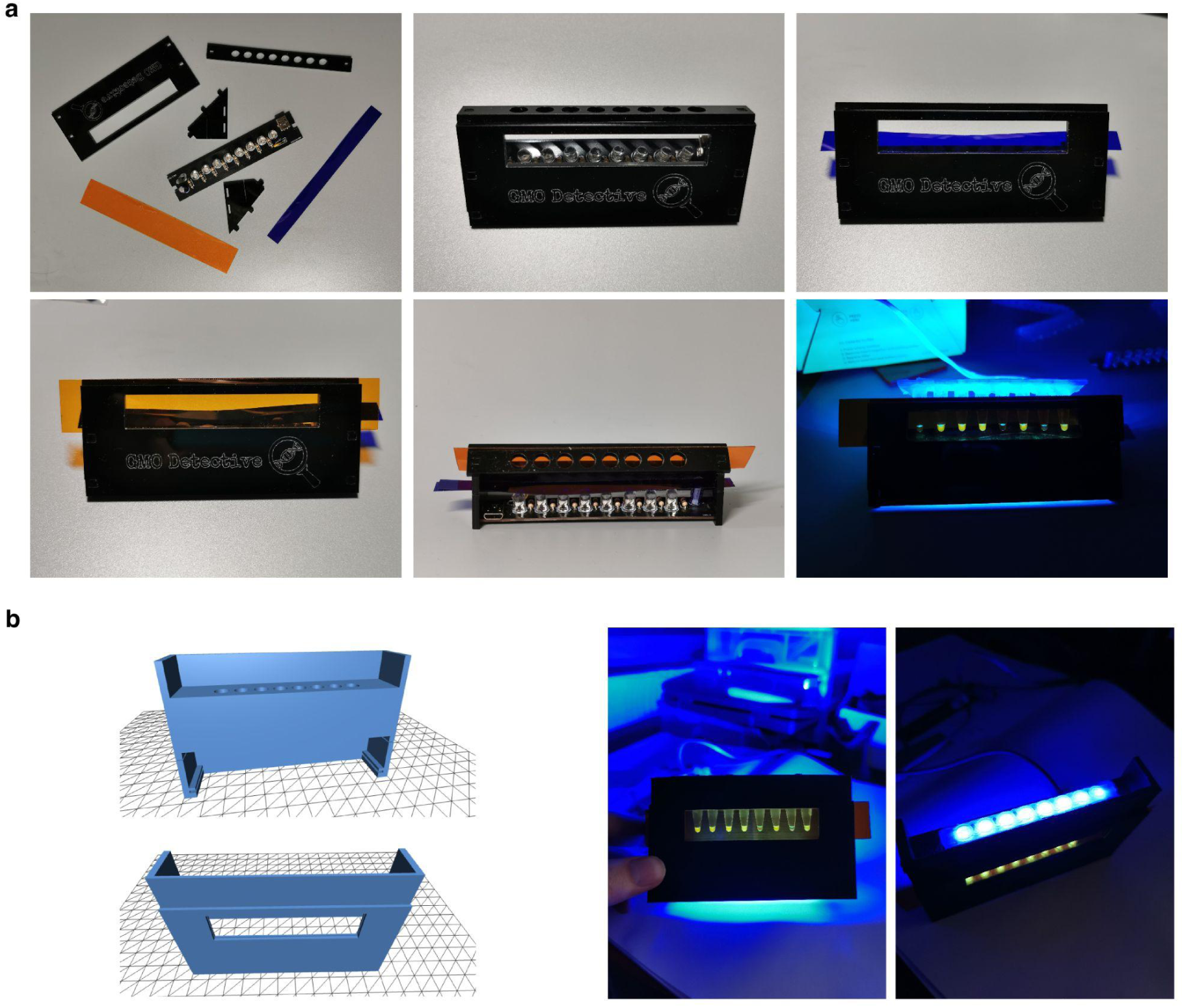
Comparison of acrylic and 3D printed GMO Detective device. a) Acrylic version of the GMO Detective device. b) 3D printed version designed in openSCAD to accommodate the same filters and PCB used in the original device. All editable files available in https://github.com/MakerLabCRI/GMODetective-Detector/blob/main/3D%20Model/STL/GMO detective3D.stl

**Figure S4:**
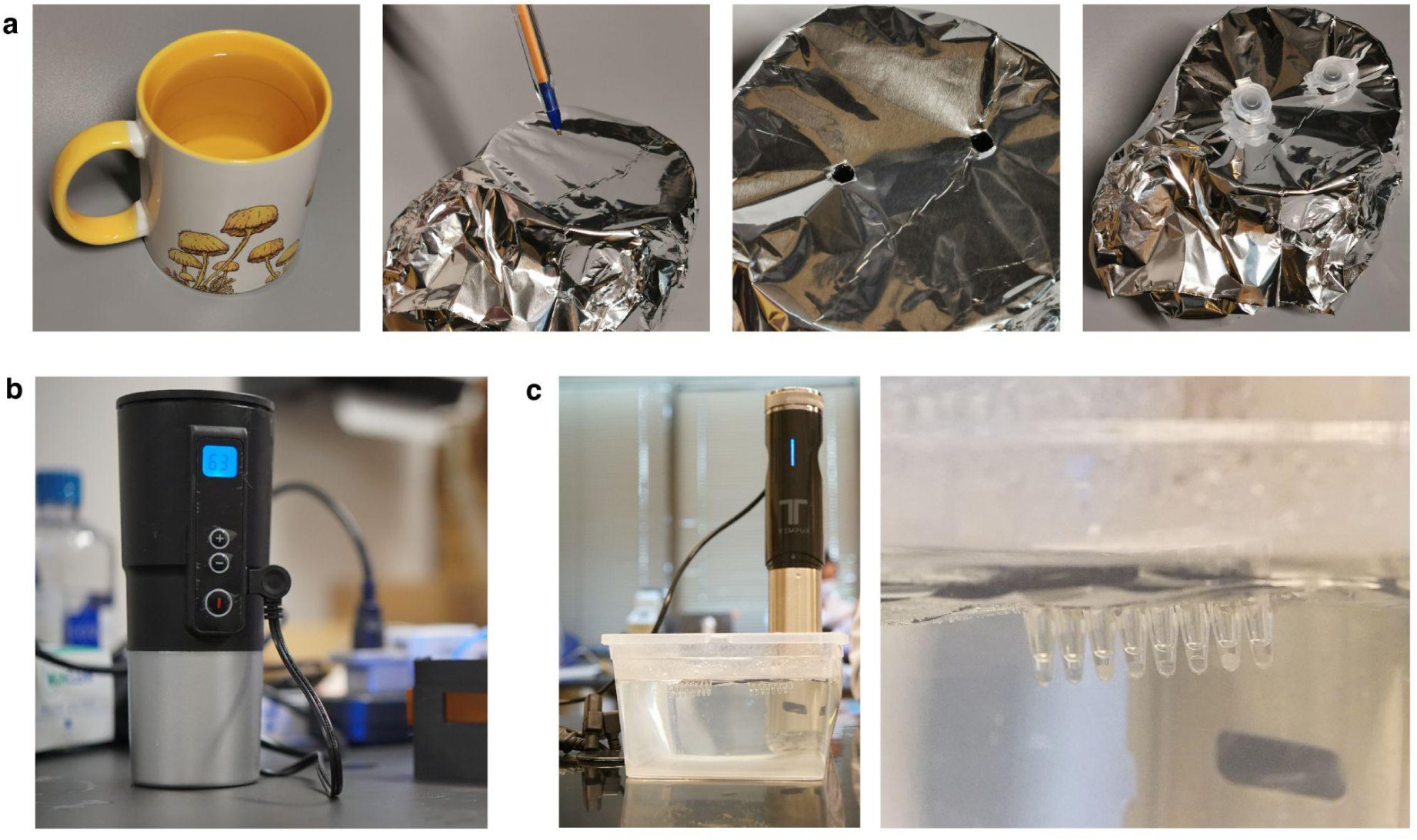
Heat incubation for sample and LAMP reactions. a) Step-by-step preparation of 100°C water incubation using aluminum foil for the initial 5 min heat treatment. b) Electronic coffee cup used for the 65°C incubation step. c) Sous vide heating device in a water bath used for the 65°C incubation step.

**Figure S5:**
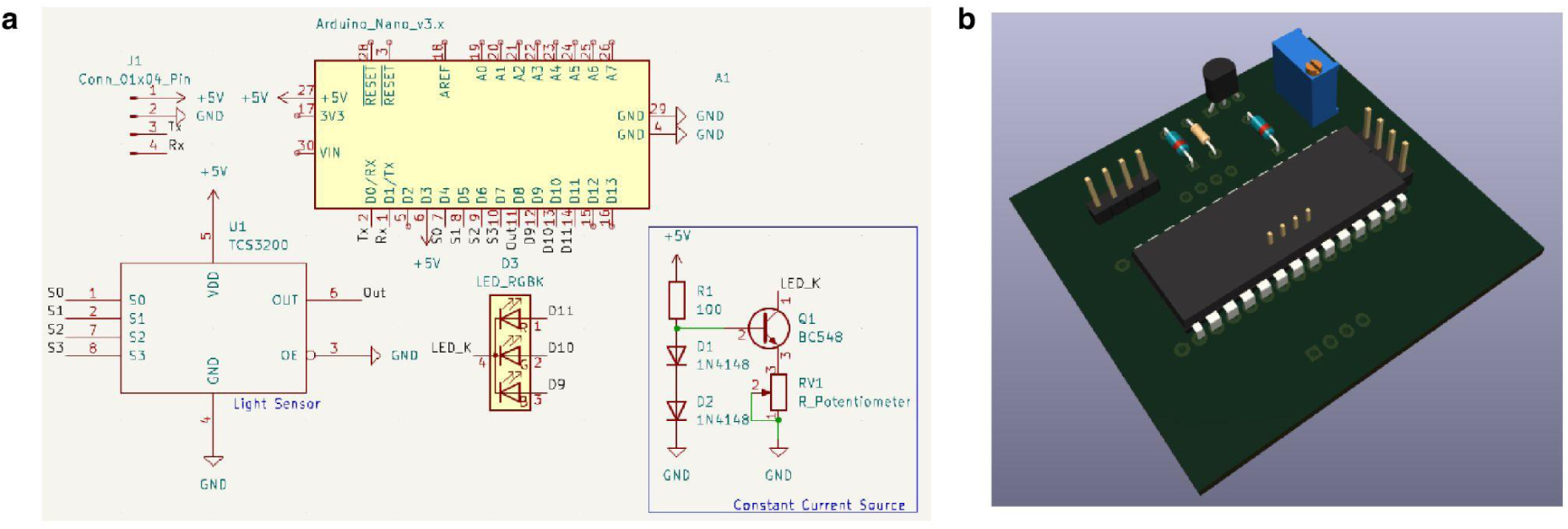
Colorimeter reGOSH. a) Schematic of the Arduino shield for the colorimeter reGOSH. b) 3D render of the colorimeter PCB. More details in https://gitlab.fcen.uncu.edu.ar/regosh/colorimetro-regosh

**Figure S6:**
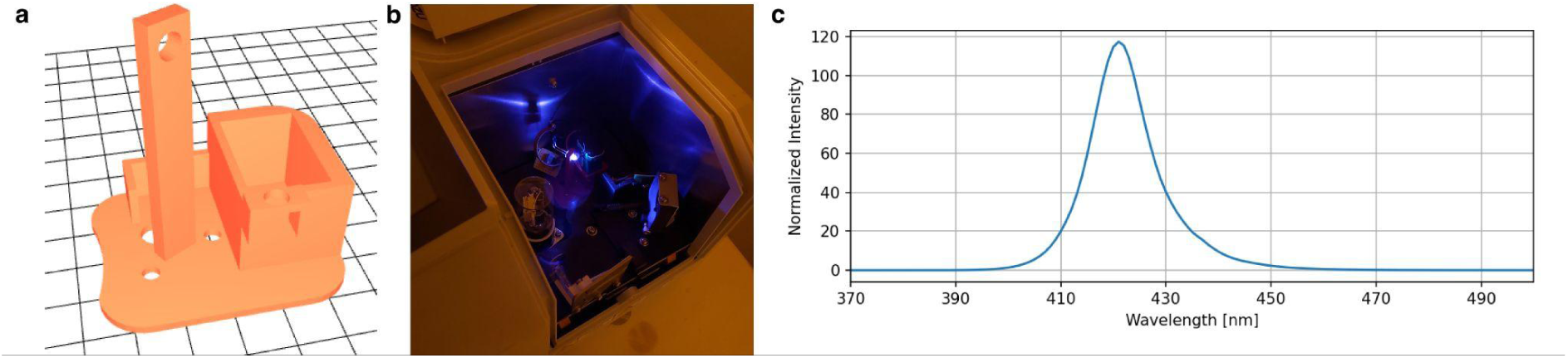
LED spectrum measurement. a) Schematic representation of the 3D printed LED adapter for the spectrometer’s light source compartment. b) UV LED placed inside the spectrometer. c) Relative intensity spectrum of the UV LED.

**Figure S7:**
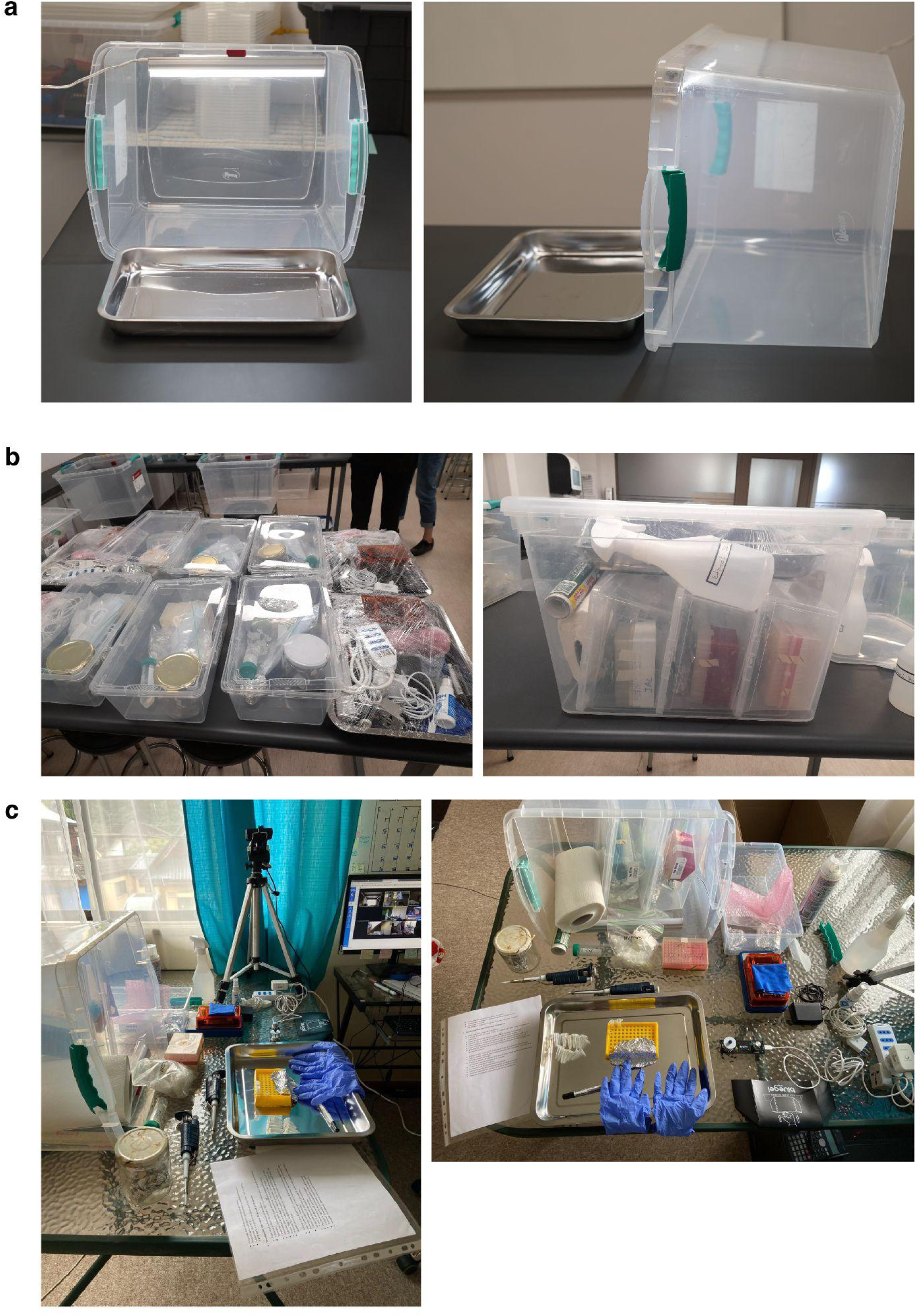
**Organization of the lab-box**. a) the box consisted of a commercial plastic storage box, which was modified to hold a light on the top via a 3D printed holder. A light metallic tray was provided to run experiments at home. The box contained three smaller plastic boxes, one per student, which can be used as a water bath for LAMP experiments. b) organization of the three smaller boxes inside the lab-in-a-box. c) The correct employment of the box was monitored by Zoom video calls (image courtesy of Josefina Lara).

## Bibliography

1. Huang A, Bryan B, Kraves S, Alvarez-Saavedra E, Stark JC. Implementing Hands-On Molecular and Synthetic Biology Education Using Cell-Free Technology. Methods Mol Biol. 2022;2433: 413–432.

2. Jung JK, Rasor BJ, Rybnicky GA, Silverman AD, Standeven J, Kuhn R, et al. At-Home, Cell-Free Synthetic Biology Education Modules for Transcriptional Regulation and Environmental Water Quality Monitoring. ACS Synth Biol. 2023;12: 2909–2921.

3. Rybnicky GA, Dixon RA, Kuhn RM, Karim AS, Jewett MC. Development of a Freeze-Dried CRISPR-Cas12 Sensor for Detecting in the Secondary Science Classroom. ACS Synth Biol. 2022;11: 835–842.

4. sobre regosh – REGOSH. [cited 1 May 2025]. Available: https://regosh.libres.cc/sobre-regosh/

5. Home - Gathering for Open Science Hardware. In: Gathering for Open Science Hardware - Gathering for Open Science Hardware [Internet]. Gathering for Open Science Hardware; 24 Feb 2016 [cited 1 May 2025]. Available: https://openhardware.science/

6. Street A. Diagnostic Development Closer to Home: The 1st Meet and Greet of the Global Frugal Diagnostic Network. In: Springer Nature [Internet]. 28 Jun 2023 [cited 1 May 2025]. Available: https://microbiologycommunity.nature.com/posts/diagnostic-development-closer-to-hom e-the-1st-meet-and-greet-of-the-global-frugal-diagnostic-network

7. Frugal Science. In: Frugal Science [Internet]. [cited 1 May 2025]. Available: https://www.frugalscience.org/

8. Hackteria. [cited 1 May 2025]. Available: https://www.hackteria.org/

9. DIYbio. In: DIYbio [Internet]. [cited 1 May 2025]. Available: https://diybio.org/

10. GLOBAL COMMUNITY BIO SUMMIT. In: GLOBAL COMMUNITY BIO SUMMIT [Internet]. [cited 1 May 2025]. Available: https://www.biosummit.org/

11. iGEM. [cited 1 May 2025]. Available: https://igem.org/

12. Biomaker.org. In: Biomaker.org [Internet]. [cited 1 May 2025]. Available: https://www.biomaker.org/

13. Website. Available: https://stanford.freegenes.org/

14. Open Bioeconomy Lab – Scientific tools for an open and equitable bioeconomy. [cited 1 May 2025]. Available: https://openbioeconomy.org/

15. Reagent Collaboration Network – a global collaboration for equitable access to biotechnology. [cited 1 May 2025]. Available: https://reclone.org/es/

16. Tahamtan A, Ardebili A. Real-time RT-PCR in COVID-19 detection: issues affecting the results. Expert Rev Mol Diagn. 2020;20: 453–454.

17. Chan JF-W, Yip CC-Y, To KK-W, Tang TH-C, Wong SC-Y, Leung K-H, et al. Improved Molecular Diagnosis of COVID-19 by the Novel, Highly Sensitive and Specific COVID-19-RdRp/Hel Real-Time Reverse Transcription-PCR Assay Validated and with Clinical Specimens. J Clin Microbiol. 2020;58. doi:10.1128/JCM.00310-20

18. Jin Y-H, Cai L, Cheng Z-S, Cheng H, Deng T, Fan Y-P, et al. A rapid advice guideline for the diagnosis and treatment of 2019 novel coronavirus (2019-nCoV) infected pneumonia (standard version). Mil Med Res. 2020;7: 4.

19. Graham TGW, Dugast-Darzacq C, Dailey GM, Nguyenla XH, Van Dis E, Esbin MN, et al. Open-source RNA extraction and RT-qPCR methods for SARS-CoV-2 detection. PLoS One. 2021;16: e0246647.

20. Cerda A, Rivera M, Armijo G, Ibarra-Henriquez C, Reyes J, Blázquez-Sánchez P, et al. An Open One-Step RT-qPCR for SARS-CoV-2 detection. PLoS One. 2024;19: e0297081.

21. Matute T, Nuñez I, Rivera M, Reyes J, Blázquez-Sánchez P, Arce A, et al. Homebrew reagents for low-cost RT-LAMP. J Biomol Tech. 2021;32: 114–120.

22. Aidelberg G, Aronoff R, Eliseeva T, Quero FJ, Vielfaure H, Codyre M, et al. Corona Detective: a simple, scalable, and robust SARS-CoV-2 detection method based on reverse transcription loop-mediated isothermal amplification. J Biomol Tech. 2021;32: 89–97.

23. PocketPCR – The thermocycler for the rest of us. [cited 1 May 2025]. Available: https://gaudi.ch/PocketPCR/

24. Reagent Collaboration Network – a global collaboration for equitable access to biotechnology. [cited 1 May 2025]. Available: https://reclone.org/es/

25. Mendoza-Rojas G, Sarabia-Vega V, Sanchez-Castro A, Tello L, Cabrera-Sosa L, Nakamoto JA, et al. A low-cost and open-source protocol to produce key enzymes for molecular detection assays. STAR Protoc. 2021;2: 100899.

26. Rivera M, Reyes J, Blazquez-Sanchez P, Ramirez Sarmiento CA. Recombinant protein expression and purification of codon-optimized Pfu-Sso7d protocol v2. 2021 [cited 1 May 2025]. Available: https://www.protocols.io/view/recombinant-protein-expression-and-purification-of-bzusp6we/v2.pdf

27. Rivera M, Reyes J, Blazquez-Sanchez P, Ramirez Sarmiento CA. Recombinant expression and purification of codon-optimized M-MLV and Mashup. 2021 [cited 1 May 2025]. Available: https://www.protocols.io/view/recombinant-expression-and-purification-of-codon-o-bsernbd6.pdf

28. González-González E, Trujillo-de SG, Lara-Mayorga IM, Martínez-Chapa SO, Alvarez MM. Portable and accurate diagnostics for COVID-19: Combined use of the miniPCR thermocycler and a well-plate reader for SARS-CoV-2 virus detection. PloS one. 2020;15. doi:10.1371/journal.pone.0237418

29. Oliveira BB, Veigas B, Baptista PV. Isothermal Amplification of Nucleic Acids: The Race for the Next “Gold Standard.” Front Sens. 2021;2: 752600.

30. Notomi T, Okayama H, Masubuchi H, Yonekawa T, Watanabe K, Amino N, et al. Loop-mediated isothermal amplification of DNA. Nucleic acids research. 2000;28. doi:10.1093/nar/28.12.e63

31. Aidelberg G. Towards democratization of nucleic acid detection. Université Paris Cité. 2021. Available: http://www.theses.fr/2021UNIP5071

32. GMO Detective – an open, affordable, reliable, easy to use, rapid, and robust kit for Gene detection by non-scientists. [cited 1 May 2025]. Available: https://gmodetective.com/

33. Ball CS, Light YK, Koh CY, Wheeler SS, Coffey LL, Meagher RJ. Quenching of Unincorporated Amplification Signal Reporters in Reverse-Transcription Loop-Mediated Isothermal Amplification Enabling Bright, Single-Step, Closed-Tube, and Multiplexed Detection of RNA Viruses. Analytical chemistry. 2016;88. doi:10.1021/acs.analchem.5b04054

34. GitHub - AlejoArav/BIO266E: Jupyter notebooks and data files for the activities carried out in the course “Biochemistry Laboratory II: Molecular Genetics.” In: GitHub [Internet]. [cited 1 May 2025]. Available: https://github.com/AlejoArav/BIO266E

35. Xu J, Wang J, Zhong Z, Su X, Yang K, Chen Z, et al. Room-temperature-storable PCR mixes for SARS-CoV-2 detection. Clinical biochemistry. 2020;84. doi:10.1016/j.clinbiochem.2020.06.013

36. Matute T, Núñez I, Federici F. Low Cost LAMP and RT-LAMP. 2021 [cited 1 May 2025]. Available: https://www.protocols.io/view/low-cost-lamp-and-rt-lamp-bsejnbcn.pdf

37. Rivera M, Cazaux S, Cerda A, Arce Medina A, Núñez I, Matute T, et al. Recombinant protein expression and purification of codon-optimized Bst-LF polymerase v1. protocols.io. ZappyLab, Inc.; 2020. doi:10.17504/protocols.io.bksrkwd6

38. Aidelberg G, Aronoff R. Freeze-drying (Lyophilization) and manufacturing of Corona Detective assay. 2020 [cited 1 May 2025]. Available: https://www.protocols.io/view/freeze-drying-lyophilization-and-manufacturing-of-bpv4mn 8w.pdf

39. GitHub - MakerLabCRI/GMODetective-Detector: DIY fluorescence detector for the GMO Detective project. In: GitHub [Internet]. [cited 1 May 2025]. Available: https://github.com/MakerLabCRI/GMODetective-Detector

40. Open source DIY kit — LED Transilluminator Build. [cited 1 May 2025]. Available: http://public.iorodeo.com/docs/led_transilluminator/

41. Nuñez I, Matute T, Herrera R, Keymer J, Marzullo T, Rudge T, et al. Low cost and open source multi-fluorescence imaging system for teaching and research in biology and bioengineering. PLOS ONE. 2017;12: e0187163.

42. Amazon.com: GEEZO 12 Oz Coffee Mug Warmer Set,Double Layer Stainless Steel Coffee Cup with Lid, 20W Wireless Induction Charging Heating Pad Up to 131°F/55°C,Perfect for Office and Home Desk (Black): Home & Kitchen. [cited 1 May 2025]. Available: https://www.amazon.com/gp/product/B08L64GBR3/ref=ppx_od_dt_b_asin_title_s00?ie=UTF8&psc=1

43. Kellner MJ, Ross JJ, Schnabl J, Dekens MPS, Matl M, Heinen R, et al. A Rapid, Highly Sensitive and Open-Access SARS-CoV-2 Detection Assay for Laboratory and Home Testing. Frontiers in molecular biosciences. 2022;9. doi:10.3389/fmolb.2022.801309

44. LEE Filters Gels. In: LEE Filters [Internet]. [cited 1 May 2025]. Available: https://leefilters.com/lighting/colour-effect-lighting-filters/

45. RGB_fluo. In: GitLab [Internet]. [cited 1 May 2025]. Available: https://gitlab.com/FernanFederici/rgb_fluor

46. Srinivasan B. Words of advice: teaching enzyme kinetics. The FEBS journal. 2021;288. doi:10.1111/febs.15537

47. Medina AA, Contreras JP, Federici F. Preparation of cell-free RNAPT7 reactions. 2017 [cited 1 May 2025]. Available: https://www.protocols.io/view/preparation-of-cell-free-rnapt7-reactions-kz2cx8e.pdf

48. Information on EC 3.2.1.23 - beta-galactosidase. [cited 1 May 2025]. Available: https://www.brenda-enzymes.org/enzyme.php?ecno=3.2.1.23#KM%20VALUE%20[mM]

49. Colorimetro reGOSH. In: GitLab [Internet]. [cited 1 May 2025]. Available: https://gitlab.fcen.uncu.edu.ar/regosh/colorimetro-regosh

50. Open Colorimeter. In: IO Rodeo [Internet]. [cited 1 May 2025]. Available: https://iorodeo.com/pages/open-colorimeter%20

51. CoSensores (Sensores Comunitarios). In: GitLab [Internet]. [cited 1 May 2025]. Available: https://gitlab.com/cosensores

52. docs/EspectroLEDs.md · main · Pablo Cremades / MultiSpect Wine Lab ·. In: GitLab [Internet]. [cited 1 May 2025]. Available: https://gitlab.fcen.uncu.edu.ar/pcremades/multispect-wine-lab/-/blob/main/docs/EspectroLEDs.md

53. Barthet MM. Teaching molecular techniques at home: Molecular biology labs that can be performed anywhere and enable hands-on learning of restriction digestion/ligation and DNA amplification. Biochem Mol Biol Educ. 2021;49: 598–604.

54. YouTube. [cited 2 May 2025]. Available: https://www.youtube.com/watch?v=b5z3K8YAUCA

55. Website. Available: https://sdgs.un.org/2030agenda

56. Chagas AM, Molloy JC, Prieto-Godino LL, Baden T. Leveraging open hardware to alleviate the burden of COVID-19 on global health systems. PLoS Biology. 2020;18: e3000730.

57. Website. Available: https://www.unesco.org/en/open-science/about

58. Bringing Open Source to the Global Lab Bench. In: Issues in Science and Technology [Internet]. 7 Feb 2022 [cited 21 May 2022]. Available: https://issues.org/open-source-science-hardware-gosh-arancio-dosemagen/

59. Arancio J. Supporting Open Science Hardware in Academia: Policy Recommendations for Science Funders and University Managers. [cited 1 May 2025]. doi:10.5281/zenodo.8030029

60. Arce A, Guzman Chavez F, Gandini C, Puig J, Matute T, Haseloff J, et al. Decentralizing Cell-Free RNA Sensing With the Use of Low-Cost Cell Extracts. Front Bioeng Biotechnol. 2021;9: 727584.

61. Bhadra S, Paik I, Torres J-A, Fadanka S, Gandini C, Akligoh H, et al. Preparation and use of cellular reagents: a low-resource molecular biology reagent platform. Current protocols. 2022;2: e387.

62. Website. Available: https://www.unesco.org/en/legal-affairs/recommendation-open-educational-resources-oer.

63. Diederich B, Müllenbroich C, Vladimirov N, Bowman R, Stirling J, Reynaud EG, et al. CAD we share? Publishing reproducible microscope hardware. Nat Methods. 2022. doi:10.1038/s41592-022-01484-5

64. GitHub - parente/nbestimate: Estimate of Public Jupyter Notebooks on. In: GitHub [Internet]. [cited 1 May 2025]. Available: https://github.com/parente/nbestimate

65. Rule A, Birmingham A, Zuniga C, Altintas I, Huang S-C, Knight R, et al. Ten simple rules for writing and sharing computational analyses in Jupyter Notebooks. PLOS Computational Biology. 2019;15: e1007007.

66. Engelberger F, Galaz-Davison P, Bravo G, Rivera M, Ramírez-Sarmiento CA. Developing and Implementing Cloud-Based Tutorials That Combine Bioinformatics Software, Interactive Coding, and Visualization Exercises for Distance Learning on Structural Bioinformatics. Journal of Chemical Education. 2021 [cited 1 May 2025]. doi:10.1021/acs.jchemed.1c00022

67. Building an Open-Source DNA Assembler Device. [cited 1 May 2025]. Available: https://ieeexplore.ieee.org/document/10019279

68. Gerhardt KP, Olson EJ, Castillo-Hair SM, Hartsough LA, Landry BP, Ekness F, et al. An open-hardware platform for optogenetics and photobiology. Scientific Reports. 2016;6: 1–13.

69. Milias-Argeitis A, Rullan M, Aoki SK, Buchmann P, Khammash M. Automated optogenetic feedback control for precise and robust regulation of gene expression and cell growth. Nature Communications. 2016;7: 1–11.

70. Quero FJ, Aidelberg G, Vielfaure H, de Kermadec YH, Cazaux S, Pandi A, et al. qByte: Open-source isothermal fluorimeter for democratizing analysis of nucleic acids, proteins and cells. bioRxiv. 2024. p. 2024.11.03.621723. doi:10.1101/2024.11.03.621723

71. Haseloff J. AirFlow microreactor. In: Hackster.io [Internet]. [cited 1 May 2025]. Available: https://www.hackster.io/jim-haseloff/airflow-microreactor-657300

72. Bhupathi M, Hegde S, Devarapu GCR, Molloy JC. MobileLAMP: A portable, low-cost, open-source device for isothermal nucleic acid amplification. bioRxiv. 2024. p. 2024.02.13.580127. doi:10.1101/2024.02.13.580127

73. Abriata LA. How Technologies Assisted Science Learning at Home During the COVID-19 Pandemic. DNA and Cell Biology. 2022;41: 19.

74. Fernandes PA, Passos Ó, Ramos MJ. Necessity is the Mother of Invention: A Remote Molecular Bioinformatics Practical Course in the COVID-19 Era. Journal of Chemical Education. 2022 [cited 1 May 2025]. doi:10.1021/acs.jchemed.1c01195

75. Biological Circuit Design — Biological Circuit Design documentation. [cited 1 May 2025]. Available: http://be150.caltech.edu.s3-website-us-west-2.amazonaws.com/2021/index.html

76. iGEM. [cited 1 May 2025]. Available: https://igem.org/

77. BioBits. [cited 1 May 2025]. Available: https://www.mybiobits.org/

78. BioBuilder Educational Foundation – Empowering the Next Generation of Scientists Through Hands-On Learning. [cited 1 May 2025]. Available: https://biobuilder.org/

79. Learning Labs. In: miniPCR bio [Internet]. [cited 8 Apr 2025]. Available: https://www.minipcr.com/products/minipcr-learning-labs/

80. The OpenFlexure project. [cited 1 May 2025]. Available: https://openflexure.org/

81. PocketPCR – The thermocycler for the rest of us. [cited 1 May 2025]. Available: https://gaudi.ch/PocketPCR/

82. Grout I. Remote Laboratories as a Means to Widen Participation in STEM Education. Education Sciences. 2017;7: 85.

83. Home - Gathering for Open Science Hardware. In: Gathering for Open Science Hardware - Gathering for Open Science Hardware [Internet]. Gathering for Open Science Hardware; 24 Feb 2016 [cited 1 May 2025]. Available: https://openhardware.science/

84. Website. Available: https://trendinafrica.org/

85. Welcome to Counter Culture Labs! In: Counter Culture Labs [Internet]. [cited 1 May 2025]. Available: https://www.counterculturelabs.org/

86. Welcome to. [cited 1 May 2025]. Available: http://www.gaudi.ch/GaudiLabs/?page_id=2

87. Molloy J. Launch of the Community Biology Biosafety Handbook. [cited 1 May 2025]. Available: https://openbioeconomy.org/blogpost/community-biology-biosafety-handbook/

88. Knowledge Hub - Manuals, Advice, Tips and Tricks. In: Bento Lab [Internet]. [cited 8 Apr 2025]. Available: https://bento.bio/resources/

89. BioBuilder. In: Carolina Knowledge Center [Internet]. 19 Apr 2023 [cited 12 Apr 2025]. Available: https://knowledge.carolina.com/our-brands/biobuilder/

90. Gogec Conference. In: Gogec [Internet]. [cited 12 Apr 2025]. Available: https://www.gogecconference.org/

91. Research use only 2019-novel coronavirus (2019-nCoV) real-time RT-PCR primers and probes. 2020 [cited 2 May 2025]. Available: https://stacks.cdc.gov/view/cdc/88834

92. Martínez FN, Medina AA, Federici F. CFE Expression and Lyophilization of β-Galactosidase (LacZ). 2023 [cited 2 May 2025]. Available: https://www.protocols.io/view/cfe-expression-and-lyophilization-of-galactosidase-cd8vs9w6.pdf

93. Cloning vector pL0R-lacZ, complete sequence - Nucleotide - NCBI. [cited 2 May 2025]. Available: https://www.ncbi.nlm.nih.gov/nuccore/1779817396

94. [No title]. [cited 2 May 2025]. Available: https://gmodetective.com/wp-content/uploads/2018/10/GMO-Detective-Wetware-V1.1.pdf?189db0&189db0

95. Open Educational Resources. 18 Jul 2022 [cited 21 Aug 2022]. Available: https://www.unesco.org/en/communication-information/open-solutions/open-educational-resources

