## Supplementary material for "Open Educational Resources for distributed hands-on teaching in molecular biology": S1 appendix

**S1 Appendix: homemade GMO Detective**

Based on the original GMO detective designed, produced and shipped from France [1], we made some modifications to primers as listed below:

**Original 35S primer set**

| Name | Sequence 5'to3' | Modification |
|---|---|---|
| 35S_FIP | aggcatcttcaacgatggccttaaaggaaggtggctcctaca | |
| 35S_BIP | tgccgacagtggtcccaaagttgaagacgtggttggaacg | 5' FAM |
| 35S_F3 | tgcccagctatctgtcactt | |
| 35S_B3 | tcccttacgtcagtggagat | |
| 35S_FLoop | tcctttatcgcaatgatg | |
| 35S_BLoop | agcatcgtgggaaaagaag | |
| 35S_BIPQ | ACTGTCGGCA | 3' Black Hole Quencher®-1 Or Iowa Black® FQ |

**Original COX1 primer set**

| Name | Sequence 5'to3' | Modification |
|---|---|---|
| COX F3 | tatgggagccgtttttgc | |
| COX B3 | aactgctaagrgcattcc | 5' FAM |
| COX FIP | atggatttgrcctaaagtttcagggcaggatttcactattgggt | |
| COX BIP | tgcatttcttagggctttcggatccrgcgtaagcatctg | |
| COX F-Loop | atgtccgaccaaagattttacc | |
| COX B-Loop | gtatgccacgtcgcattcc | |
| COX FIPQ | YCAAATCCAT | 3' Black Hole Quencher®-1 orIowa Black® FQ |

**Modified 35S primer set (modifications labeled in yellow)**

| Name | Sequence 5'to3' | Modification |
|---|---|---|
| 35SZ-F3 | aagatgcctctgccgaca | |
| 35SZ-B3 | cagcgtgtcctctccaaat | |
| 35SZ-FIP | acgtggttggaacgtcttcttcccaaagatggaccccca | |
| 35SZ-BIP | atctccactgacgtaagggatgatagaggaagggtcttgcga | 5' FAM |
| 35SZ-LF | tccacgatgctcctcgtg | |
| 35SZ-LB | acgcacaatcccactatcct | |
| 35SZ-BIP-Q | cagtggagat | 3' Iowa Black FQ |

**Modified COX1 primer set (modifications labeled in yellow)**

| Name | Sequence 5'to3' | Modification |
|---|---|---|
| COX-F3 | tatgggagccgtttttgc | |
| COX-B3 | aactgctaagRgcattcc | |
| COX-FIP | atggatttgRcctaaagtttcagggcaggatttcactattgggt | 5' Texas Red |
| COX-BIP | tgcatttcttagggctttcggatccRgcgtaagcatctg | |
| COX-LF | atgtccgaccaaagattttacc | |
| COX-LB | gtatgccacgtcgcattcc | |
| COX-FIP-Q | YCAAATCCAT | 3' Iowa Black FQ |

*FAM*: Fluorescein
*Iowa Black FQ*: Fluorescence quencher in the FAM range.

QUASR probe diagram used:

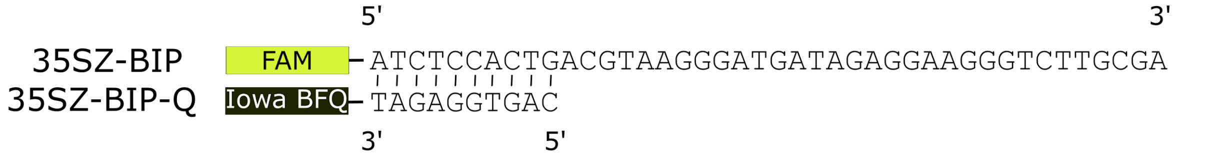


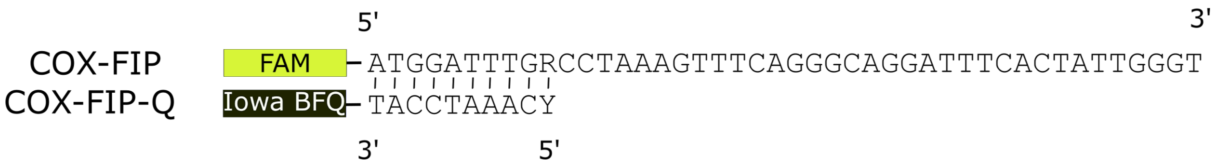


**Annealing of LAMP primers for S.tuberosum mitochondrial cox1 gene** [2]:

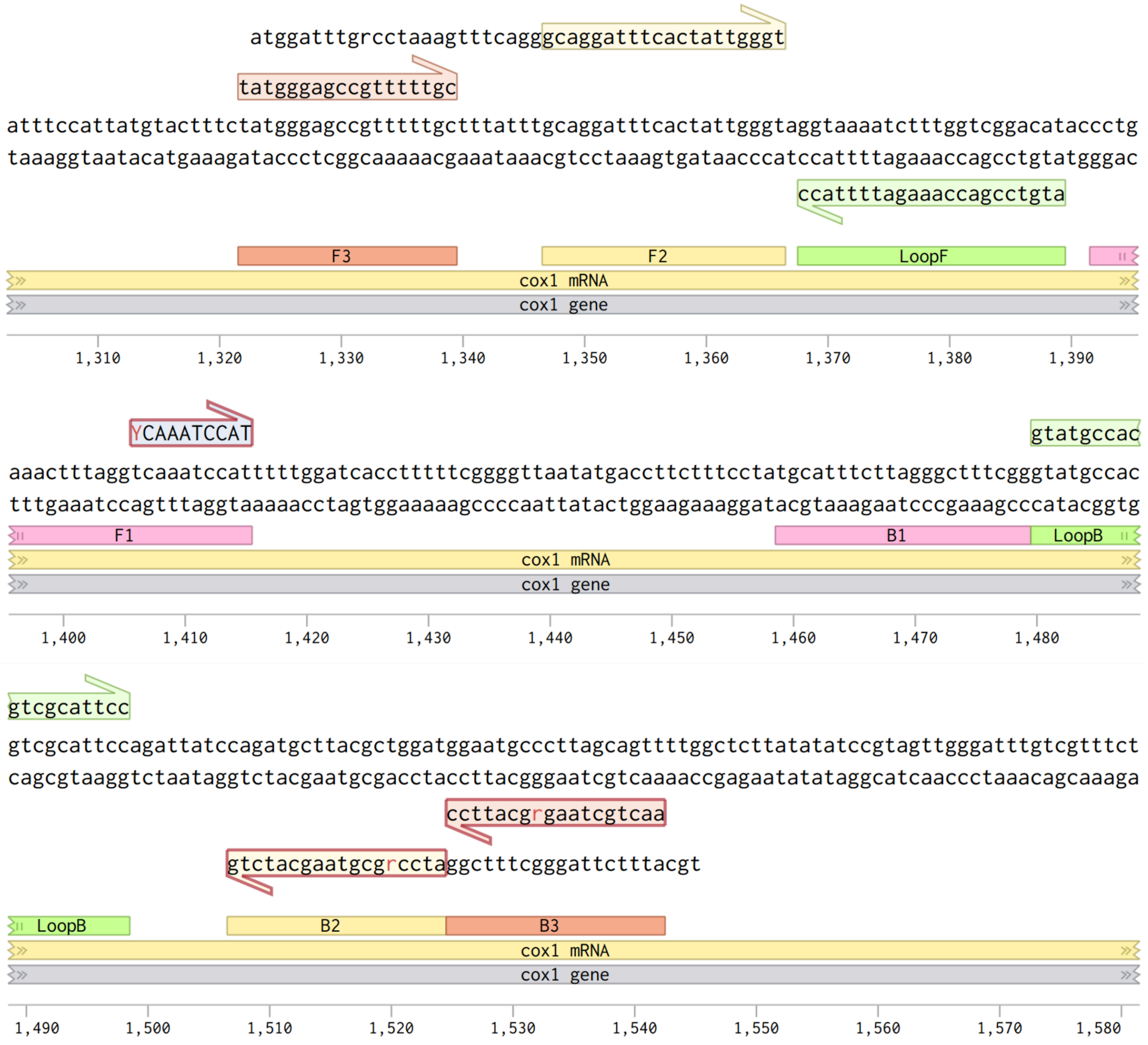

### **Annealing of LAMP primers for 35S CaMV** [3]

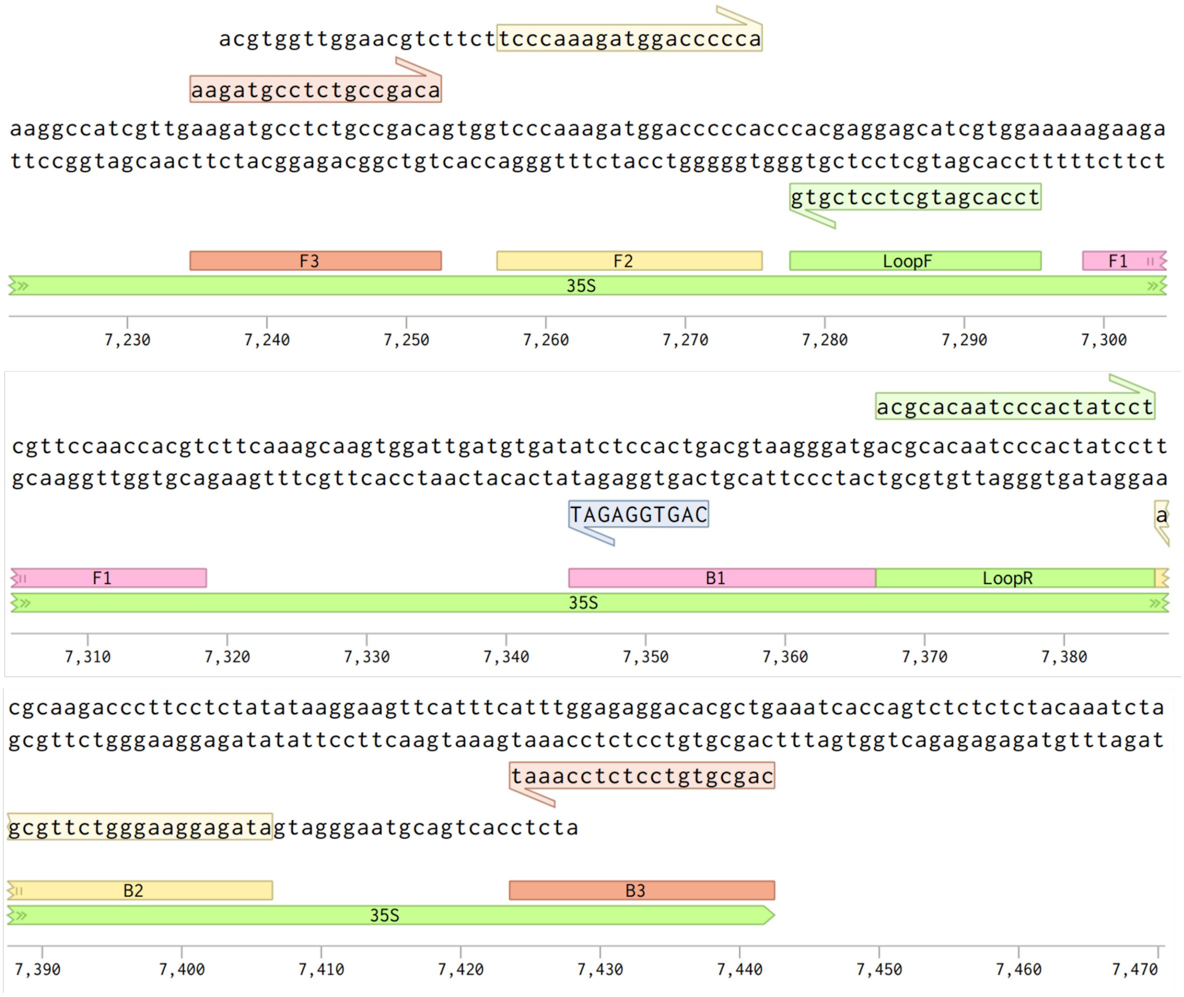


#### **Primer quantities and purification methods at ordering**

| **Name** | **Sequence** | **Scale** | **Purification** | | |
|---|---|---|---|---|---|
| COX-F3 | tatgggagccgtttttgc | 100nm | STD | 100 nmole DNA oligo | Standard desalting |
| COX-B3 | aactgctaagRgcattcc | 100nm | STD | 100 nmole DNA oligo | Standard desalting |
| COX-FIP | /5TexRd-XN/atggatttgR cctaaagtttcagggcaggattt cactattgggt | 250nm | HPLC | 250 nmole DNA oligo | **HPLC purification** |

| | | | | | |
|---|---|---|---|---|---|
| COX-BIP | tgcatttcttagggctttcggatc cRgcgtaagcatctg | 250nm | STD | 250 nmole DNA oligo | Standard desalting |
| COX-LF | atgtccgaccaaagattttacc | 100nm | STD | 100 nmole DNA oligo | Standard desalting |
| COX-LB | gtatgccacgtcgcattcc | 100nm | STD | 100 nmole DNA oligo | Standard desalting |
| COX-FIP-Q | YCAAATCCAT/3IAbRQ Sp/ | 250nm | HPLC | 250 nmole DNA oligo | **HPLC purification** |
| 35SZ-F3 | aagatgcctctgccgaca | 100nm | STD | 100 nmole DNA oligo | Standard desalting |
| 35SZ-B3 | cagcgtgtcctctccaaat | 100nm | STD | 100 nmole DNA oligo | Standard desalting |
| 35SZ-FIP | acgtggttggaacgtcttcttcc caaagatggaccccca | 250nm | STD | 250 nmole DNA oligo | Standard desalting |
| 35SZ-BIP | /56-FAM/atctccactgacgt aagggatgatagaggaaggg tcttgcga | 250nm | STD | 250 nmole DNA oligo | Standard desalting |
| 35SZ-LF | tccacgatgctcctcgtg | 100nm | STD | 100 nmole DNA oligo | Standard desalting |
| 35SZ-LB | acgcacaatcccactatcct | 100nm | STD | 100 nmole DNA oligo | Standard desalting |
| 35SZ-BIP-Q | C AGT GGA GAT/3IABkFQ/ | 250nm | HPLC | 250 nmole DNA oligo | **HPLC purification** |

Thermocycler

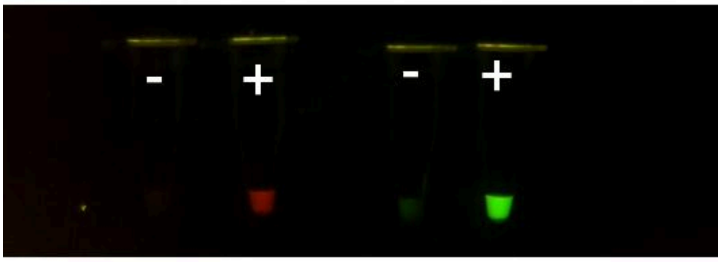


3D printer hot bed

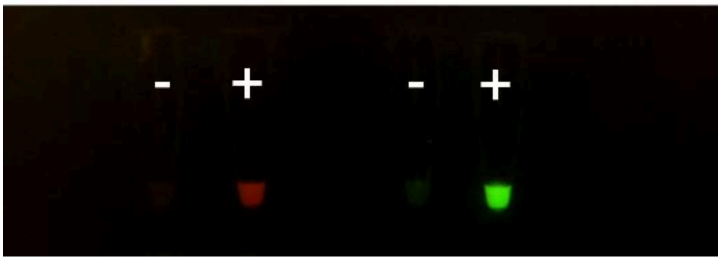

Comparison of Texas Red COX1 (left) and FAM 35S (right) QUASR LAMP reactions run on a thermocycler (top) and 3D printer hot bed (bottom). The thermocycler was run at at 63°C while the heated bed was run at 65°C. Homemade enzymes and buffers were used in both. The photo was taken after 60 min of incubation in a FluoPi [4](at 200 ISO, 760000 µs exposure time) .


1. [No title]. [cited 22 Feb 2024]. Available: https://gmodetective.com/wp-content/uploads/2018/10/GMO-Detective-Wetware-V1.1.pdf?189db0&189db0

2. COX1_X83206 · Benchling. [cited 22 Feb 2024]. Available: https://benchling.com/s/seq-jsZWJHYQhWAf3DYHAdUo

3. 35s_V00141 · Benchling. [cited 22 Feb 2024]. Available: https://benchling.com/s/seq-McxTSERY7eYYw9cOe7u9

4. Nuñez I, Matute T, Herrera R, Keymer J, Marzullo T, Rudge T, et al. Low cost and open source multi-fluorescence imaging system for teaching and research in biology and bioengineering. PLoS One. 2017;12: e0187163.
