## Supplementary material for "Open Educational Resources for distributed hands-on teaching in molecular biology": S2 appendix

### S2 Appendix: RGBfluor design and operation instructions

#### RGBfluor printed circuit board (PCB)

The PCB was designed to be compatible with the commercial NodeMCU v3 "Lolin" board based on the ESP8266 microcontroller. The PCB has a shield format and has 9 digital controlled WS2812B RGB LEDs.

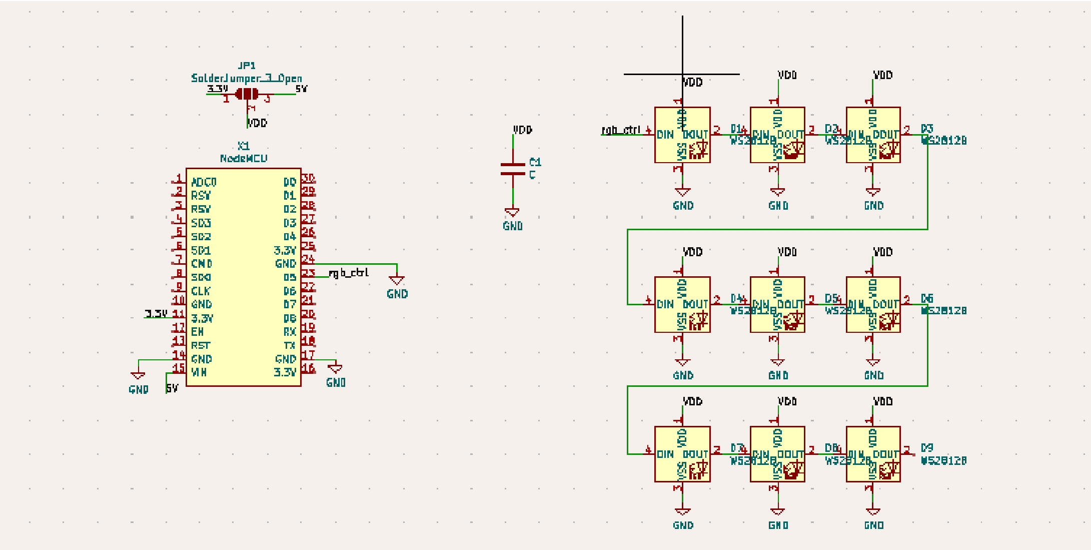

Schematic of the Lolin (left) and the LEDs of RGBfluor PCB (right).

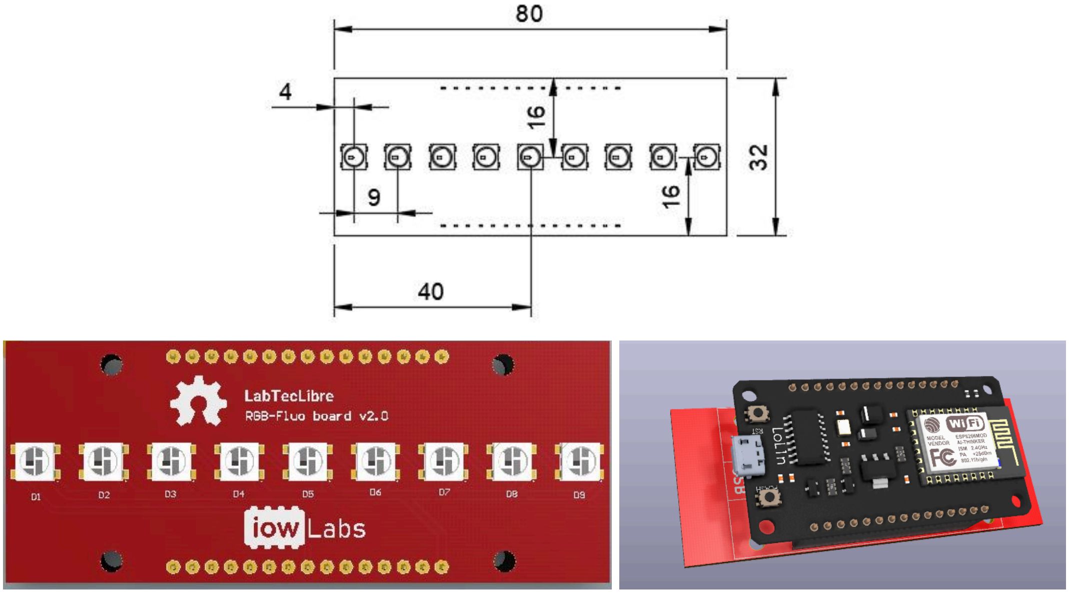


Schematic representation of the physical dimension and spatial arrangement of RGB LEDS in the RGBfluor PCB.

Bill of materials for the construction of the circuit is:

| Item | Name | Quantity | Part number LCSC | Note |
|---|---|---|---|---|
| Capacitor | C1 | 1 | C77386 | MLCC SMD 0603 1uF. |

| | | | | |
|---|---|---|---|---|
| RGB LED | D1-D9 | 9 | C114586 | WS2812B |
| Header | J1 | 2 | C2897378 | FH, 1 row x 15 pos. pitch 2.54. |

The gerber files to replicate the board are available in the repository https://gitlab.com/FernanFederici/rgb_fluor under the path PCB/Gerber_Files. The board is simple to assemble and the components used can be assembled by hand soldering.

##### Firmware and software

The firmware was developed on platformIO with the Arduino framework. It implements a local software access point and mounts a local server on the ESP8266 board, which can be accessed from any device connected to the generated network. When the device is powered, the microcontroller will generate a local network with a specific name (by default RGB-Fluo 0X), which can be configured by programming the firmware.
The assembled network is discoverable by any device with the ability to connect via Wi-Fi protocol: PC, tablet, smartphone, and has no password. Once entered the network we can access to the web application by entering the following IP
http://192.168.4.1/

The code uses the following libraries to work (all of them available from the official Arduino library manager)

- ESP8266 WiFi
- me-no-dev/ESPAsyncTCP
- me-no-dev/ESPAsyncWebServer
- LittleFS
- fastled/FastLED

The ESP8266WiFi, ESPAsyncTCP and ESP Async WebServer libraries are used to manage WiFi communication, implement the access point and set up the local server. The LittleFS library implements a memory management (File system) for the microcontroller, allowing the html and css files from the web page to be stored in the flash memory of the board. The FastLED library allows you to control RGB LEDs. These LEDs can be controlled individually, however for the purposes of this application they are all controlled at the same time.

##### Compiling the code

To upload the code to the device, some preliminary steps are necessary. You have to mount an image in the memory of the microcontroller with the files index.html and syles.ccs that contain the web page found in the path

./firmware/data

You have to run the commands.

pio runs -t buildfs pio runs -t uploadfs

Once the image is uploaded we can load the firmware through the commands

Compile: run pio Load: pio run --target load
