## Supplementary material for "Open Educational Resources for distributed hands-on teaching in molecular biology": S3 appendix

S3 Appendix: RGBfluor student guide

### RGB Fluor

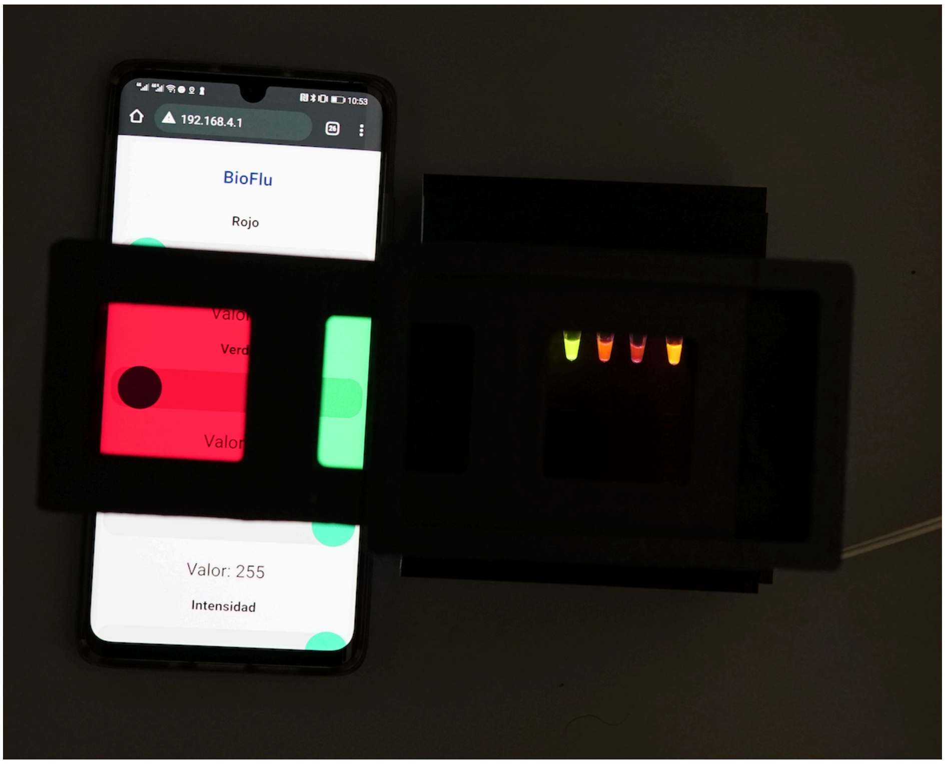


This document corresponds to the support material for the fluorescence practical of the Bio266e course.

Authors: Fernan Federici, Anibal Arce, Wladimir Araya, Domingo Gallardo, Isaac Nuñez, Tamara Matute.

This material is licensed under Creative Commons BY-SA: This license allows reusers to distribute, remix, adapt and build upon the material in any medium or format, as long as attribution is given to the creator. The license allows commercial use. If you remix, adapt or build on the material, you must license the modified material under identical terms (https://creativecommons.org/licenses/by-sa/4.0/deed.es).

**Practical learning objectives:**

- To understand the organization of an optical system for the detection of a fluorescent signal; particularly the compatibility between illumination wavelength, optical properties of the emission filter and excitation/emission spectrum of the fluorescent molecules.

**Task:**

Applying the concepts learned in class on fluorescence microscopy, we ask you to take a photograph with your mobile phone of:

1- The best setup to photograph the protein **sfGFP**
2- The best setup to photograph the protein **mScarlet-I**
3- The best setup to photograph both proteins:**sfGFP and mScarlet-I simultaneously**.

(The idea is that the photograph has a dark background to increase the range between the signal and the background. That is, a photo where GFP is seen on an all-green background, for example, is not useful).

You must consider compatibility between:

1. Excitation/emission spectrum of fluorescent molecules.
2. Optical properties of the emission filter
3. Lighting Wavelength

**1) Excitation/emission spectrum of fluorescent molecules.**

You can find the profiles of both fluorescent proteins in these links:
sfGFP: https://www.fpbase.org/protein/superfolder-gfp/
mScarlet-I: https://www.fpbase.org/protein/mscarlet-i/

| Name | $\lambda_{ex}$ | $\lambda_{em}$ | Stokes |
|---|---|---|---|
| Superfolder GFP | 485 | 510 | 25 |
| mScarlet-I | 569 | 593 | 24 |

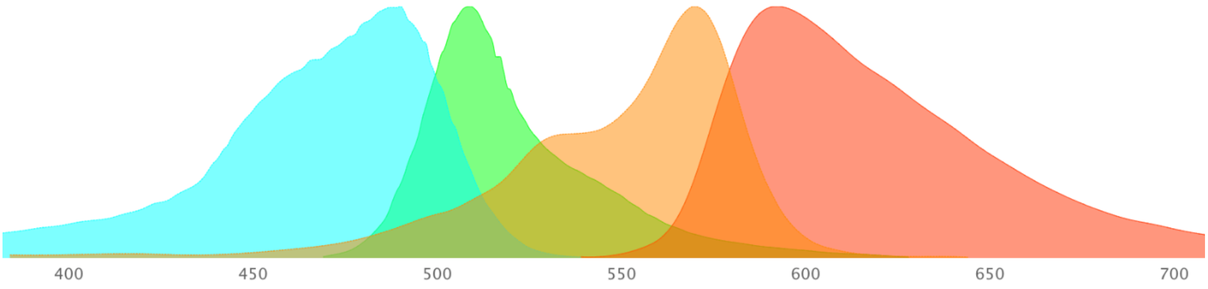


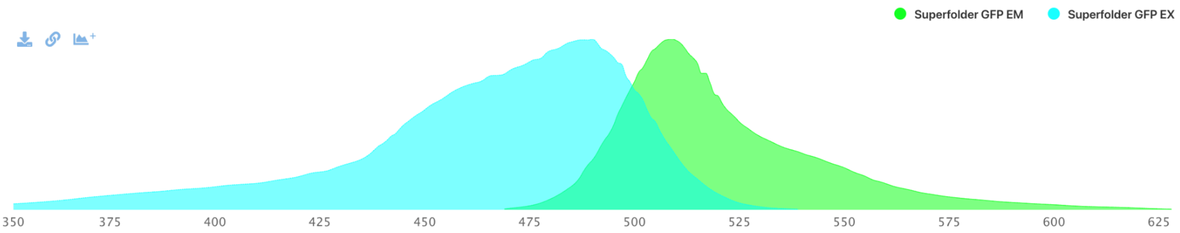


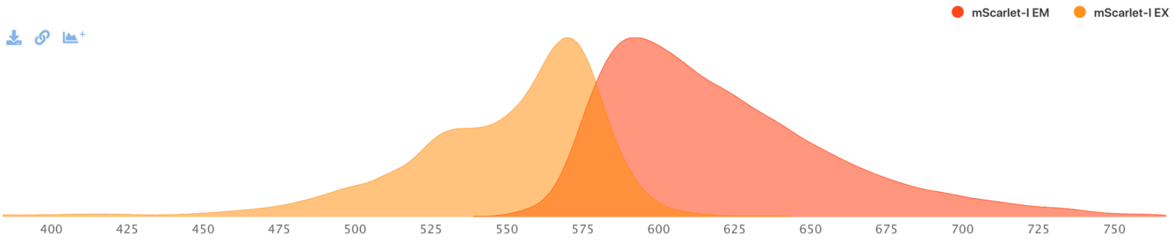


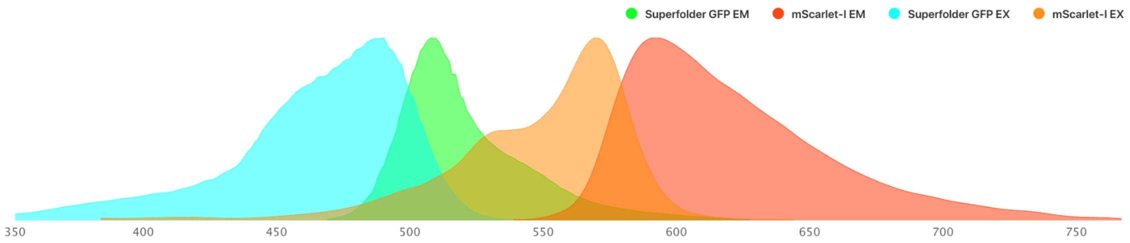


Attribute Comparison

| Name | $\lambda_{ex}$ | $\lambda_{em}$ | Stokes | EC | QY | Brightness | pKa | Aggregation | Maturation | Lifetime | kDa |
|---|---|---|---|---|---|---|---|---|---|---|---|
| Superfolder GFP | 485 | 510 | 25 | 83,300 | 0.65 | 54.15 | | wd | 13.6 | | 26.78 |
| mScarlet-I | 569 | 593 | 24 | 104,000 | 0.54 | 56.16 | 5.4 | m | 36.0 | 3.1 | 26.36 |

#### 2) optical properties of emission filter

105: http://www.leefilters.com/lighting/colour-details.html#105
113: http://www.leefilters.com/lighting/colour-details.html#113&filter=cf
122: http://www.leefilters.com/lighting/colour-details.html#122

If you click on the links and move the cursor over the spectrogram you will be able to see the exact values:

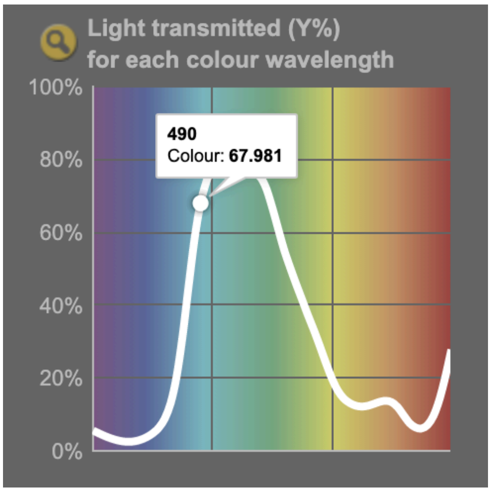


These filters have very different optical properties that will allow you to observe sfGFP and mScarlet-I with different results.

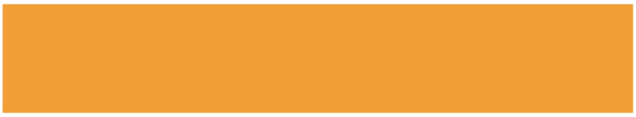

**105 Orange**

View on leefilters.com / Find a Dealer

Good for light entertainment and functions. Creates a good fire effect when used with 106 or 104.

| | Source C 6774K | Tungsten 3200K |
|---|---|---|
| Transmission Y | 41.3% | 50% |
| x | 0.563 | 0.59 |
| y | 0.428 | 0.406 |
| Absorption | 0.38 | 0.3 |

**Light transmitted (Y%)** for each colour wavelength (nm)

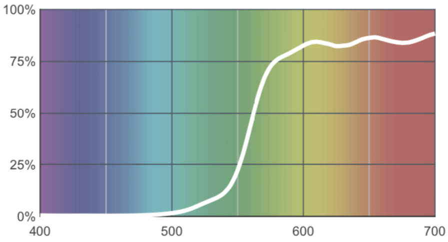

#### 113 Magenta

View on leefilters.com / Find a Dealer

Very strong - used carefully for small areas on set.

| | Source C 6774K | Tungsten 3200K |
|---|---|---|
| Transmission Y | 10.9% | 14.7% |
| x | 0.563 | 0.662 |
| y | 0.217 | 0.29 |
| Absorption | 0.96 | 0.83 |

**Light transmitted (Y%)** for each colour wavelength (nm)

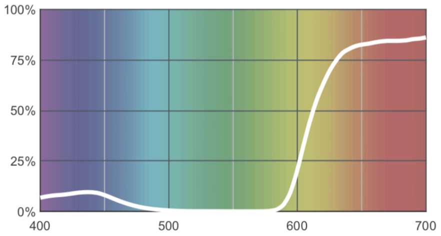


#### 122 Fern Green

View on leefilters.com / Find a Dealer

Good for cycloramas and creates a great mood effect.

| | Source C 6774K | Tungsten 3200K |
|---|---|---|
| Transmission Y | 51.5% | 45.7% |
| x | 0.234 | 0.336 |
| y | 0.543 | 0.549 |
| Absorption | 0.29 | 0.34 |

**Light transmitted (Y%)** for each colour wavelength (nm)

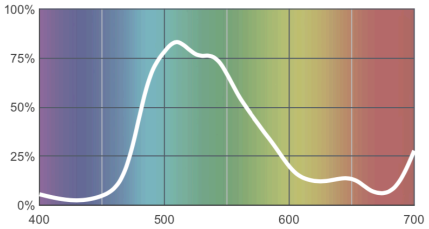


#### 3) lighting wavelength

The lighting will be obtained through RGB LED lights, which have lighting Blue (465-467 nm), verde (522-525 nm) and red (620-625nm).

**Instructions for use:**

This device, called RGBfluor, is made up of a top part that houses the three lee filters (described above) and a bottom part that contains the lighting plate and fluorescent tubes. The modules at the bottom do not need to be manipulated. The interaction with the device will occur via 1) sliding the module with the three filters and 2) activating the corresponding lighting with your phone.

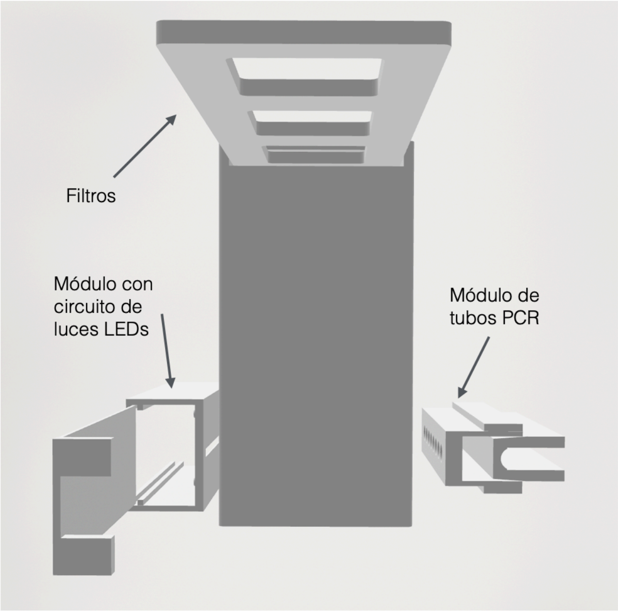


To control the LED lights, you must connect your phone to the WIFI signal called BioFluo1, 2, 3...depending on the device you have. Once your phone is connected, you must open a web page with the following IP:**192.168.4.1** and the following control interface will open:

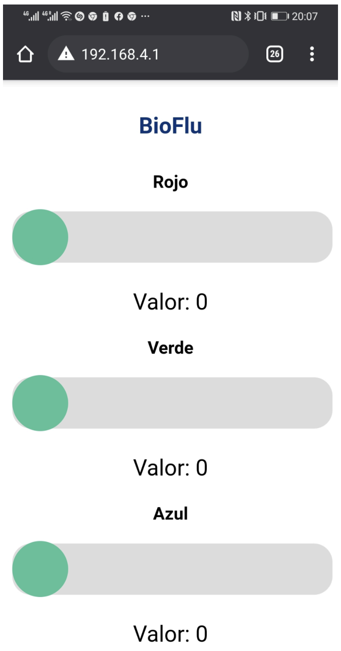

From this interface you can control the color of the excitation light of your samples.

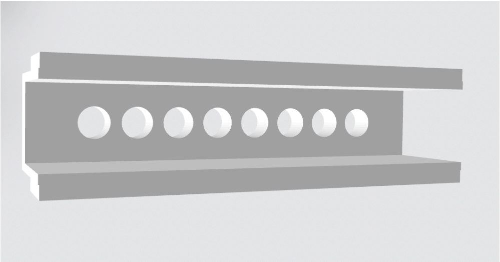

A) PCR tube container

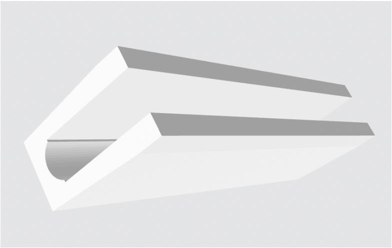

B) PCR tube holding cap

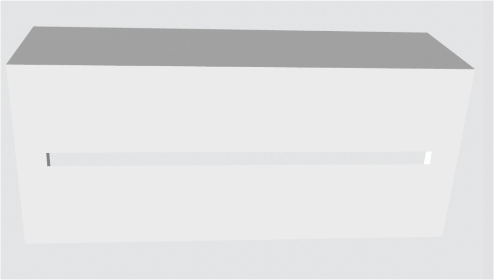

C) PCB circuit container with LED lights

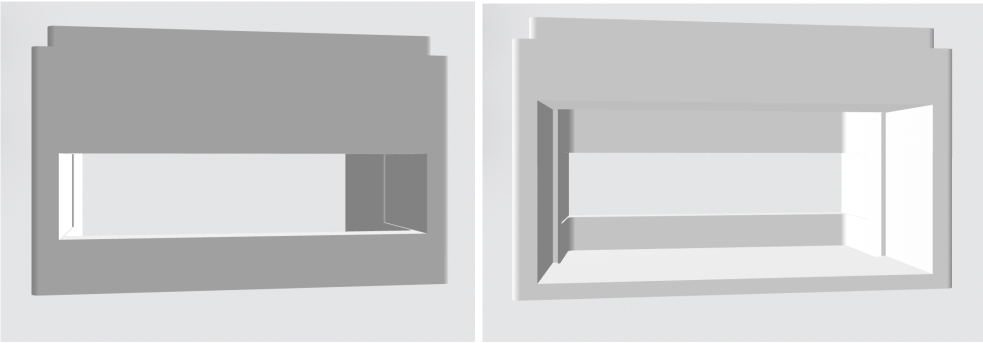

D) Base of the RGB fluo, showing the two windows where, on the left, the PCR tube container is inserted (A and B) and the one on the right showing where the PCB container (C) is inserted.

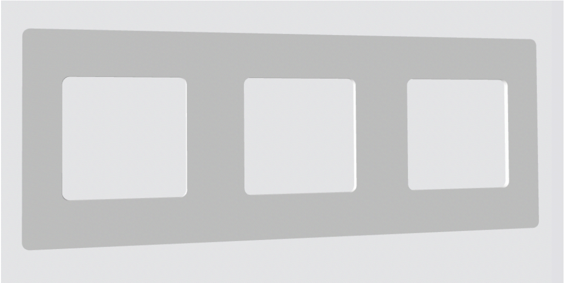

E) Container of the three filters

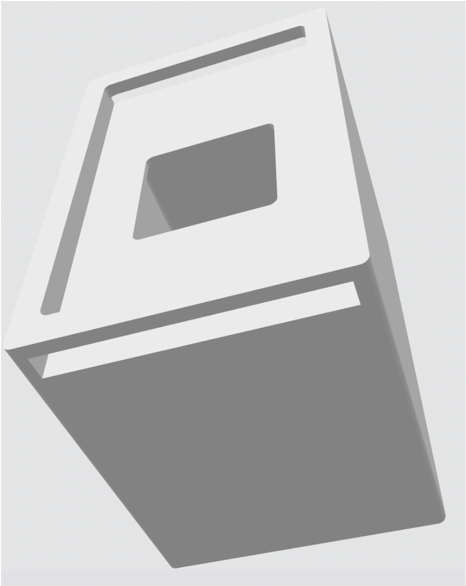

F) Top of the device

**Links of interest:**

- Complete list of fluorescent proteins:https://docs.google.com/spreadsheets/d/1RFIFflIkxFzspQ8SHJcLsHKuf8X3u9uaLJn0-eQRJk8/edit#gid=354984915 (very useful in the future!)
- The best portal on fluorescent proteins:https://www.fpbase.org/
- Digital repository with design files of the open hardware device to be used:https://gitlab.com/FernanFederici/rgb_fluor
- Protein production protocol in “cell-free” systems:https://www.protocols.io/view/preparation-of-cell-free-rnapt7-reactions-kz2cx8e

(Please leave us your comments where you think the document needs improvements.)
