## Supplementary material for "Open Educational Resources for distributed hands-on teaching in molecular biology": S4 appendix

### S4 Appendix: Student`s guide fo practical on enzyme kinetics

In this practical exercise, we will use β-galactosidase enzyme from E. *coli*, an enzyme that degrades β-galactoside disaccharides to monosaccharides, including lactose to glucose and galactose. The substrate used in this activity will be a synthetic substrate called **o-nitrophenyl-β-D-galactopyranoside (oNPGal)**. oNPGal is used because the assay method is simpler than using the natural substrate lactose, as one of the reaction products, **o-nitrophenol (oNP)**, absorbs at 420 nm and can be quantified by absorbance (e.g., with a colorimeter, spectrophotometer or by image analysis).

During this practical experiment, a characterization of *E. coli* β-galactosidase enzyme produced by a cell-free transcription-translation reaction will be carried out. The lysate containing the enzyme has been lyophilized for its transport at room temperature. In the first step, students must hydrate the enzyme with 1X PBS buffer to then quantify its activity against different substrate concentrations in order to characterize its kinetic parameters (Vmax and Km). Calibration curves of the product oNP will be used to determine the amount of product produced in the reactions.

#### 1. Calibration curve for the quantification of the reaction product, o-nitrophenol (oNP)

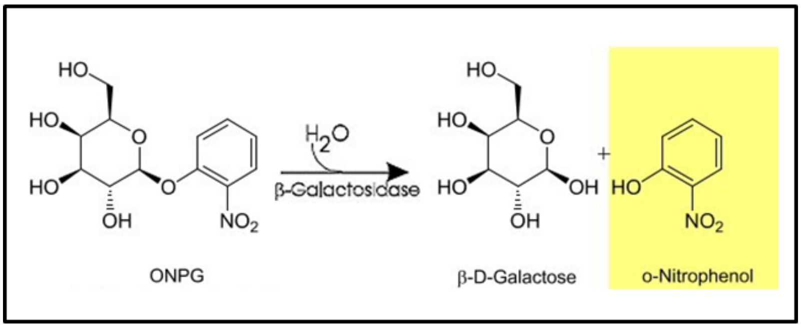

A series of oNP dilutions was previously prepared by the teaching assistants as indicated in the table below from a freshly prepared stock of oNP and using 1X PBS buffer. This curve represents the oNP calibration curve, conducted under the same buffer conditions as the reactions that will be performed to measure β-galactosidase activity. This curve will be used to interpolate the absorbance values obtained experimentally, to transform these values into units of concentration of the reaction product.

Each student must calculate the μM concentration of oNP (and fill out the last column of the table). Also, each student should measure the absorbance at 420 nm of each dilution using any of the different methods of this laboratory (spectrophotometer, colorimeter or quality images for posterior processing) and fill out the Absorbance at 420 nm column of the table.

| o-nitrophenol 10 mM (µl) | Water (µl) | PBS pH 7.4 (µl) | $Na_2CO_3$ 1M pH 10 (µl) | Absorbance at 420 nm | [oNP] µM |
|---|---|---|---|---|---|
| 0 | 400 | 100 | 500 | | |
| 1 | 399 | 100 | 500 | | |
| 3 | 397 | 100 | 500 | | |
| 5 | 395 | 100 | 500 | | |
| 10 | 390 | 100 | 500 | | |
| 15 | 385 | 100 | 500 | | |
| 20 | 380 | 100 | 500 | | |
| 25 | 375 | 100 | 500 | | |
| 30 | 370 | 100 | 500 | | |
| 45 | 355 | 100 | 500 | | |
| 60 | 340 | 100 | 500 | | |

Note: The total volume of the reaction mixture plus sodium carbonate ($Na_2CO_3$) is constant and equal to 1 ml. Absorbances should be read from the most diluted to the most concentrated ($Na_2CO_3$ is an alkaline buffer, which will halt enzymatic activity).

**2. To obtain the saturation curve for β-galactosidase, follow these steps:**

1. Rehydration of lyophilized enzyme:

Perform a quick spin-down to ensure the enzyme (should look like a white powder) is at the bottom of the tube and will not be lost when opening the cap. Add 1 ml of saline buffer (PBS) 1X to the tube containing the lyophilized enzyme. Vortex for 40 seconds to ensure homogeneous resuspension of the enzyme. Keep on ice.

2. Preparation of substrate dilutions

Prepare 9 substrate dilutions from the concentrated oNPGal stock (3 mg/ml), following the table below (including a substrate-free blank). Use water, substrate, and phosphate buffer as indicated in the table below. Remember that enzyme addition will start the reaction, so enzyme will be added as the last reagent, when all tubes are ready and timing starts. When adding the enzyme to the tubes, invert the tubes 10 times to initiate the reaction.

3. Incubation

Incubate the tubes for 13 minutes at 37°C to allow the reaction to occur.

4. Reaction termination

Add 0.5 ml of $Na_2CO_3$ 1M pH 12 to each tube to stop the reaction. Invert the tubes (closed) 5-10 times to mix well.

5. Determination of product concentration:

Measure absorbance at 420 nm of the product reaction by the same methods used for the calibration curve. Use the previously obtained calibration curve to infer the product concentration in each sample. Fill the table bellow with the absorbance values.

| Water (µl) | oNPGal 3 mg/ml (µl) | PBS pH 7.4 (µl) | Resuspended enzyme (µl) | $Na_2CO_3$ 1M pH 12 (µl) | Absorbance 420 nm |
|---|---|---|---|---|---|
| 300 | 0 | 100 | 100 | 500 | |
| 295 | 5 | 100 | 100 | 500 | |
| 294 | 6 | 100 | 100 | 500 | |
| 292 | 8 | 100 | 100 | 500 | |
| 287 | 13 | 100 | 100 | 500 | |
| 275 | 25 | 100 | 100 | 500 | |
| 250 | 50 | 100 | 100 | 500 | |
| 200 | 100 | 100 | 100 | 500 | |
| 175 | 125 | 100 | 100 | 500 | |

#### 3. Estimation of Km y Vmáx

1. Prepare a table with the following data:

| Substrate concentration µM | Absorbance 420 nm | [Product] µM | Initial velocity (Vo) µM/min | 1/[Substrate] (1/µM) | 1/Vo (1/min) |
|---|---|---|---|---|---|

2. Represent the data using the Lineweaver Burk (double reciprocal plot). The x-axis is the reciprocal of the substrate concentration, or 1 / [S], and the y-axis is the reciprocal of the reaction velocity, or 1 / V. Interception in the x axes represents 1/Km; slope represents Km/Vmax; interception with the y axis represents 1/Vmax
