## Supplementary material for "Open Educational Resources for distributed hands-on teaching in molecular biology": S5 appendix

### S5 Appendix: Box usage instructions

#### Receiving the box

When you receive the box, be careful and check it is the one corresponding to the practical that you must carry out (LAMP or RT-PCR) and student group. You can verify this information using the label on the box, which identifies what type of practical it corresponds to, the group number and the name of the students who should receive it. If you have any problem please contact the teaching team immediately.

You should keep the box in a cool place where it is not exposed to direct sunlight and **DON'T OPEN** it until the moment of carrying-out the practical.

#### What is inside the box?

You received a large box containing 3 individual smaller boxes inside and a metal tray. On the metal tray, there are elements that are common for the use of the 3 students, please, manipulate them very carefully.

The elements and reagents found inside the small boxes are for **personal use**. The material inside each small box is in sterile conditions and meant for each student's particular use. **ONLY OPEN THE SMALL BOX THAT CORRESPONDS TO YOU.**

Before sending the box to the next student, be sure to carefully clean and store each common item following the instructions in the manual.

#### Sterilization box

When you open the box, the first thing you should do is take out the gloves and put them on. Later, you will find a bottle that contains 70% ethanol and that will allow you to sterilize the large box **inside and out,** and also the small ones **only on the outside**. Apply as much 70% ethanol as you deem necessary on the surface of the boxes and use absorbent paper to distribute it over the surface. Let the ethanol dry by evaporation.

Once the practical activity is finished and after having completed the section **"Cleaning and storage of material"** found at the end of this manual, be sure to spray the entire surface of the box with 70% ethanol before storing it inside the box, following the instructions previously described.

#### The box as a workstation.

The large box will become each student's own workstation for the duration of the practical. Therefore, the first thing you should do is place it on a stable surface, avoiding irregular and slippery surfaces, in order to avoid accidents. Once you have established your workplace, you must carefully remove the tray and one small work box (the other two small boxes can remain inside the large box). Place the large box horizontally, install the LED light on the corresponding bracket and connect it to the power source using the extension cord provided inside the box.

Position the tray at the front of the box so that it is illuminated by the LED light. You must develop the practical inside the tray, so that if a material spill occurs, it does not affect your home and is easy to clean.

**Biosecurity Standards**

You will carry out a practical activity at home, but you must take all the corresponding safety measures as if you were working in a laboratory, even being even **more cautious**. Therefore, you should, if possible, carry out the practical activity in an environment free of small children and pets.

Outfit:

- You should wear long pants, closed-toed shoes and avoid wearing loose clothing.
- Those students who have long hair must wear it up, ensuring that their face is completely uncovered. Those who have bangs should ensure that they do not fall over the forehead.
- You should not use pendants or rings.
- Avoid wearing contact lenses.
- The use of gloves and mask is mandatory.

Work conduct:

- Prior to the practical activity, you must have read and studied the corresponding protocol.
- **DON'T** handle objects in your home with gloves on, to avoid contamination of your living space or work materials.
- **DON'T** use gloves to handle your mobile device. If you need to use it to show something to your co-workers or to take a photograph, you must remove your gloves.
- **DON'T** touch your face with gloves on.
- **DON'T** eat or drink during the practical activity.
- You must follow the protocol delivered for each practical activity, without making any modifications to the detailed instructions.

**IMPORTANT:** Since we are in a pandemic, you will be asked to wear a mask during the practical activity (the mask is provided inside your individual work box). This measure will prevent work material from being contaminated in the probable case of SARS-CoV-2. If you present symptoms associated with the disease caused by SARS-CoV-2, we ask you to urgently notify the teaching staff.

**Materials inside the box**

Each box has all the elements and reagents necessary to carry out the practical experiences, you **MUST NOT** use any utensils from your home.. Below you will find a list of the common and personal materials and reagents in each box.

**LAMP box**

Common materials:

- Micropipettes:
  - P200.
  - P20.
- 70% ethanol.
- Plastic base
- Power-strip.
- Beaker.
- Water Heater or Thermometer.
- Printed circuit board (*PCB*) with LEDs and acrylics.
- USB cable - microUSB.
- Tube rack (2).
- Fine tip marker.

Personal materials (small box):

- Tip box.
- Bottle with tubes.
- Food coloring.
- Reaction strip.
- Problem sample (corn).
- Plant sample: Stems (2).
- Metallic paper.
- *Parafilm*.
- Reagents:
  - Nuclease-free $H_2O$ (H).
  - Reaction buffer (R).
- Waste bag.

**RT-PCR box**

Common materials:

- Micropipettes:
  - P200.
  - P20.
- 70% ethanol.
- Plastic base
- Power strip.
- LED light.
- PocketPCR thermal cycler.
- USB-C adapter.
- 2A charger.
- blueGel electrophoresis chamber™.
- Spray ClearView™ (and its cloth).
- Tube rack.
- Fine tip marker.

Personal materials:

- Tip box.
- Bottle with tubes.
- Food coloring.
- Reagents (in parentheses is the label that you will see on the tube):
  - Nuclease-free $H_2O$ (H).
  - 5X reaction buffer (B).
  - dNTPs (N).
  - DTT (DTT).
  - Primers (4).
    - Viral Forward (FN).
    - Viral Reverse (RN).
    - Plant Forward (FA).
    - Plant Reverse (RA).
  - Enzyme mix (E).
  - Genetic material (4).
    - Viral RNA (RV).
    - Viral cDNA (DV).
    - Plant RNA (RM)
    - Plant cDNA (DM).
  - 6X Load Buffer (BC).
  - Molecular weight standard 50bp (50).
  - Positive reaction controls (2).
    - Viral (+V)
    - Plant (+M)
- Mineral oil (AC).
- 2% agarose gel.
- 50mL TAE 1X.
- Waste bag

**What to do with the waste?**

During the practical instance you will produce a lot of waste plastic material (mainly tips and tubes), which you must store inside the zip-lock bag that you will find in its individual box. The waste bag corresponds to the same bag where the reagents are located. For this purpose, remove ALL tubes from the bag, put them in their rack, and begin using the bag as waste storage. Before adding waste to the bag, make sure each tube is properly closed to prevent spillage.

Once the practical instance is over, you must seal well the bag that will contain **ALL** of the plastic waste, spray it with 70% ethanol and leave it inside the individual box. The teaching team will be responsible for properly disposing of all waste produced during the practical activity when the boxes are collected after the third student.

### Cleaning and storage of material

Store all material and waste in your personal box. Common materials must all remain inside the metal tray. Use the list of materials and reagents above to check that you do not forget any materials. The cleaning of the specific materials for each box can be found in the specific manuals for each of them.

Once all the materials are on the tray, wrap them with plastic wrap to prevent them from moving from the tray. Leave the remaining plastic wrap inside the box and close it. Now it is ready to send to the next student/back to the teaching team.

### References


Environment Cleaning and Disinfection Protocol. (2020).*Ministry of Health*, *19*, 1. https://www.minsal.cl/wp-content/uploads/2020/03/PROTOCOLO-DE-LIMPIEZA-Y-DESINFECCIÓN-DE-AMBIENTES-COVID-19.pdf

*Gob.cl - coronavirus*. (n.d.). Retrieved October 15, 2020, from https://www.gob.cl/coronavirus/?gclid=CjwKCAjw5p_8BRBUEiwAPpJO6-wOiAB38BIWpvB99-OLjE7pWEZYHSGBGJMBu4d26xUomb2CNbDJThoCbbcQAvD_BwE
