## Supplementary material for "Open Educational Resources for distributed hands-on teaching in molecular biology": S5 appendix

### S6 Appendix: Micropipetting instructions

Aim: Learn how to manipulate micropipettes correctly.

Before carrying out any practical activity, we must ensure that you know how to use micropipettes correctly and understand their volume range. This tutorial will teach you the basics of using micropipettes and will end with an activity checkup.

Before starting the corresponding LAMP or RT-PCR practical activity, you will need to upload a photograph of the pipetting practical to the task box (set up for this activity) on the CANVAS platform, this has the objective of verifying that you learned the correct handling of this delicate instrument.

#### 1. Parts of a micropipette.

- Plunger (*Push button for Plunger*): part of the micropipette that, when pressed or released, will allow a volume of liquid to be taken and released. **IMPORTANT**: The plunger has two stops, the stop to use depends on the activity being carried out, and will be explained in detail below.
- Volume control wheel (*Thumb Wheel*): allows you to adjust the volume that the micropipette will take. **IMPORTANT**: each micropipette has a volume range in which it is functional, remember **NO** to use a volume outside that range (the micropipette loses calibration).
- Volume window (*Volumeter*): indicates the adjusted volume that the micropipette will take.
- Ejector button (*Tip Ejector Button*): button that, when pressed, will remove the tip of the micropipette through the movement of the ejector.
- Tip support (*Barrel Tip Holder*): place where the tip is adjusted to suck the desired volume.
- Tip (*Tip*): place where the volume of working liquid will be stored. **IMPORTANT**: NEVER use a micropipette without a tip, the liquid can enter the micropipette, contaminate it and cause it to decalibrate.

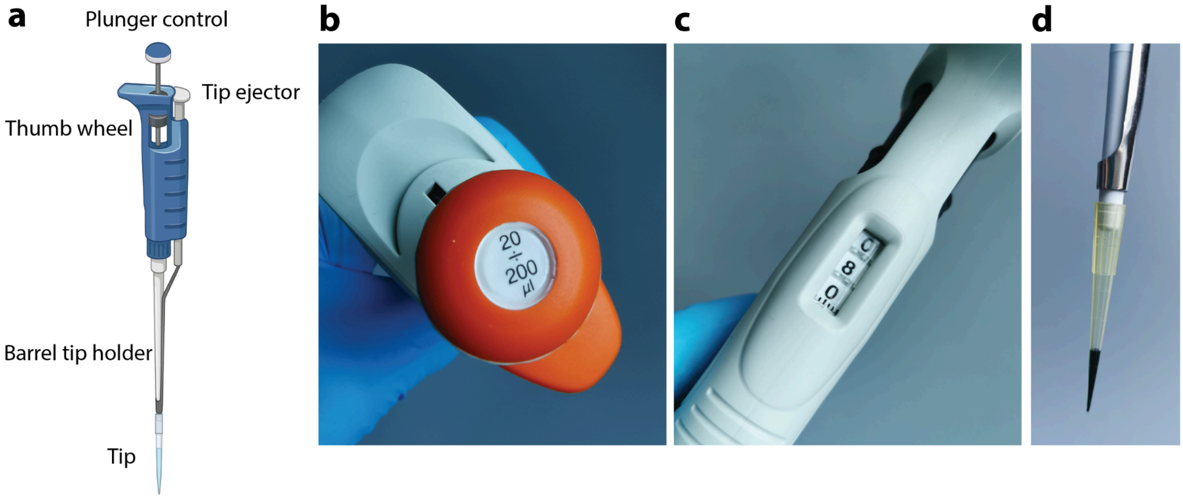


**Figure 1: Parts of a micropipette.** a) Schematic representation of a micropipette. b) Volume working range, here between 20 and 200 µL. c) volume window. d) tip containing a correct volume of liquid without bubbles.

#### 2. Types of micropipettes.

The micropipette to use in the practicals will depend on the required volume. There are different micropipettes that work in limited ranges of volumes and require different tips depending on their size.

In this practical experience, you will have two micropipettes called: P200 and P20. The name of the micropipette is given by the maximum working volume that can be used with them:

- P200: working volume between 20 and 200µL (in some cases 50 to 200µL).
- P20: working volume between 2 and 20µL.

You will have a box of tips, which are suitable for P200 and P20. The tips should be changed each time a volume of a different liquid is used to avoid contamination problems between different vials.

The accuracy of the micropipette depends on its calibration, this must be done regularly (approximately every year). For greater accuracy when taking a volume with the micropipette, you should always choose the micropipette whose maximum allowable volume is closest to this volume. For example, if you want to take 20µl, it is preferable to use the P20 rather than the P200, this is because there is less space between the surface of the liquid and the plunger of the micropipette.

#### 3. How to use a micropipette.

The proper way to hold a micropipette can be seen in Figure 2. This posture is comfortable for right- and left-handed people. The plunger is pressed with the thumb, while the rest of the fingers are in charge of holding the micropipette.

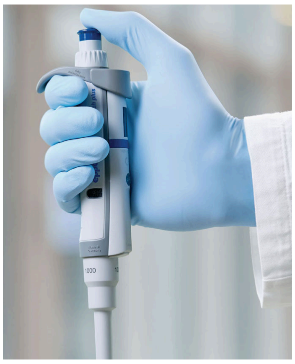

**Figure 2: correct micropipette holding.**

To suck volume for direct pipetting:

1. Adjust the workload with the wheel that allows you to control the volume.
2. Take one tip of appropriate size, verifying that it fits well in the tip support.
3. Press the plunger with the thumb up to the first stop.
4. Hold down until first stop (Figure 3) and immerse the tip in the liquid you want to suck. **CONSIDERATIONS**: Try to keep the pipette as vertical as possible or at a maximum angle of 40°. The tip of the pipette should be immersed a few millimeters into the liquid. DO NOT touch the bottom of the tube with the tip, as this prevents suction.
5. Slowly release the plunger to suck the liquid until it is no longer necessary to press it. **CONSIDERATIONS**: If the bottom of the tip does not contain liquid, but air, perform the procedure again. Air at the bottom of the tip indicates that it did not take in the proper volume.

b. To release volume:

1. Take the micropipette to the tube/paper on which the liquid will be deposited.
2. Touch the wall of the tube and slowly press the plunger until you reach the first stop, and then to the second stop (Figure 3).
3. **DO NOT** release the plunger until the micropipette moves from where it released the liquid (otherwise it will suck the liquid again).
4. Throw away the tip that has just been used in the corresponding waste, pressing the

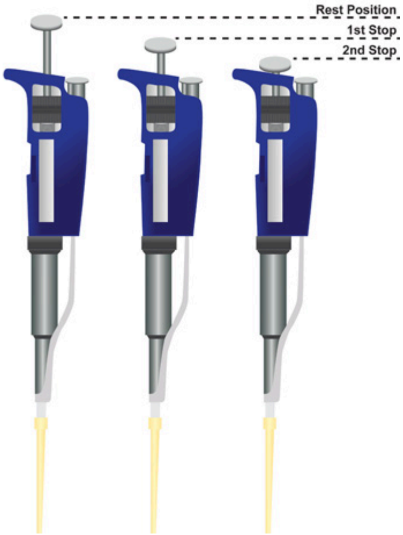


ejector button.

**Figure 3: Plunger positions for taking and releasing liquid.**

**4. Practical activity.**

Now that you know the parts of a micropipette and the theory of how to suck and release a volume of liquid, it's time to put your knowledge into practice with a small activity.

The activity consists of carrying out two serial dilutions of a food coloring.

Dilution 1:

1. Label five 0.2mL tubes with the numbers 1 to 5.
2. In a total volume of 100µL, make a 1/5 dilution of the food coloring in tube #1.
3. Perform a 1/5 dilution of tube No. 1 for tube No. 2, obtaining a final volume of 100 µL.
4. Repeat the procedure until tube No. 5.

Dilution 2:

1. Label 4 0.2mL tubes with the letters A to D.
2. In a total volume of 100µL, make a 1/10 dilution of the food coloring in tube A.
3. Perform a 1/10 dilution of tube A for tube B obtaining a final volume of 100µL.
4. Repeat the procedure until tube D.

Arrange the tubes of both dilutions together according to the color gradient, write down the pattern of numbers and letters obtained according to this order and take a photograph like the one seen in Figure 4. The pattern obtained must match the one delivered by the equipment. teacher so that you can continue with the practical activity.

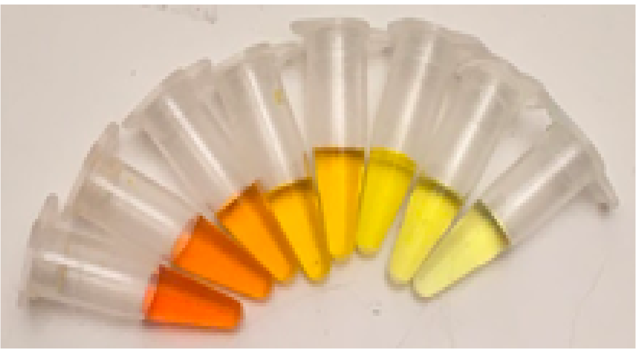

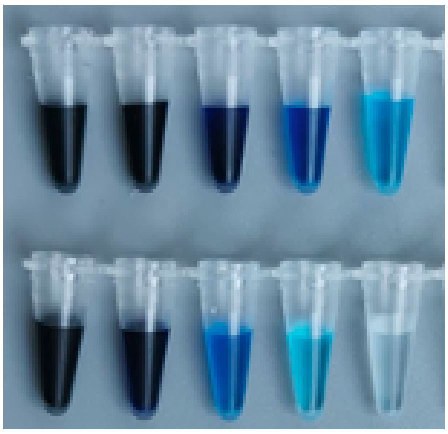

**Figure 4: Serial dilution example.**

#### References


Ewald, K. (2015). *Impact of pippetting techniques on precision and accuracy*. Userguide. https://handling-solutions.eppendorf.com/fileadmin/Community/Calibration/Userguide_020_Reference-family_Research-family_Research-pro_Impact-of-pipetting-_eng.pdf

MiniPCR. (2019). *Micropipetting 101-Micropipette Mastery Activity*. https://www.minipcr.com/wp-content/uploads/miniPCR-Learning-Labs-Micropipetting-101-240919-Micropipetting-Mastery.pdf
